# Metastable neural dynamics underlies cognitive performance across multiple behavioural paradigms

**DOI:** 10.1101/657569

**Authors:** Thomas H. Alderson, Arun L.W. Bokde, J.A.Scott. Kelso, Liam Maguire, Damien Coyle

## Abstract

Despite resting state networks being associated with a variety of cognitive abilities, it remains unclear how these local areas act in concert to express particular cognitive operations. Theoretical and empirical accounts indicate that large-scale resting state networks reconcile dual tendencies toward integration and segregation by operating in a metastable regime of their coordination dynamics. One proposal is that metastability confers important behavioural qualities by dynamically binding distributed local areas into large-scale neurocognitive entities. We tested this hypothesis by analysing fMRI data in a large cohort of healthy individuals (N=566) and comparing the metastability of the brain’s large-scale resting network architecture at rest and during the performance of several tasks. Task-based reasoning was principally characterised by high metastability in cognitive control networks and low metastability in sensory processing areas. Although metastability between resting state networks increased during task performance, cognitive ability was more closely linked to spontaneous activity. High metastability in the intrinsic connectivity of cognitive control networks was linked to novel problem solving (or fluid intelligence) but was less important in tasks relying on prior experience (or crystallised intelligence). Crucially, subjects with resting architectures similar or ‘pre-configured’ to a task-general arrangement demonstrated superior cognitive performance. Taken together, our findings support a critical linkage between the spontaneous metastability of the large-scale networks of the cerebral cortex and cognition.

## Introduction

The brains of subjects at ‘cognitive rest’ display circumscribed patterns of neural activity or resting state networks (Fox et al., 2005; Fox and Raichle, 2007) that broadly overlap with task-based activations (Smith et al., 2009; Cole et al., 2014a). Somehow these large-scale networks of the brain rearrange themselves on a fixed anatomical structure to support internal processes relevant to cognition (Lewis et al., 2009; Sadaghiani and Kleinschmidt, 2013; Bola and Sabel, 2015; Braun et al., 2015; Spadone et al., 2015; Cohen and D’Esposito, 2016; Cohen, 2018). One proposal is that neuronal assemblies are dynamically bound into coherent coordinative structures known as neurocognitive networks (Bressler and Kelso, 2001, 2016). The concept of the neurocognitive network represents an important compromise between two antagonistic viewpoints: the first, localisation, which holds that complex cognitive functions are localised to specific regions of the brain, the second, globalism, which posits that complex functions are distributed and arise through global coordination (McIntosh, 1999, 2000, 2004, 2007; Bressler and Mcintosh, 2007). From the neurocognitive network perspective, the brain’s tendencies toward integration and segregation are simultaneously realised. Local areas are permitted to express their intrinsic functionality yet also couple together and coordinate globally. Cognition, in this context, is the real-time expression of distributed local areas whose states of mutual coordination are adjusted dynamically over time (Bressler and Tognoli, 2006). An important challenge is to understand how these local areas become dynamically linked in the execution of particular cognitive operations, and equally, how these patterns of dynamic connectivity evolve over time (Cabral et al., 2017; Gonzalez-Castillo and Bandettini, 2017).

The coordination of neurocognitive networks appears to arise from a dynamic regime that balances counteracting tendencies toward integration and segregation (Tononi et al., 1994, 1998; Shanahan, 2010; Sporns, 2013; Tognoli and Kelso, 2014a). Empirical and theoretical accounts indicate that the brain derives this behaviour from its identity as a complex dynamical system operating in the metastable regime of its coordination dynamics (Kelso, 1995, 2012; Kelso and Tognoli, 2007; Tognoli and Kelso, 2009, 2014b; Shine et al., 2016b). The concept of metastability represents an important theoretical solution to the requirement that local areas operate independently yet also combine and behave synergistically (Kelso and Tognoli, 2007; Kelso, 2012; Tognoli and Kelso, 2014b). Metastability is important too as an observable phenomenon, furnishing a dynamical explanation for how large-scale brain regions coordinate their activity in space and time to support cortical function (Bressler and Kelso, 2001, 2016; Jirsa and McIntosh, 2007; Kelso, 2008; Tognoli and Kelso, 2009). In the language of dynamical system theory, metastability refers to a coupled or collective oscillatory activity which falls outside its equilibrium state for dwell times that depend on distance from equilibrium. A concrete example of a metastable dynamical system is the ‘winnerless competition’ (Rabinovich et al., 2006, 2008), however, metastable phenomena may emerge from a variety of underlying mechanisms (where certain conditions are satisfied) and it is in this broader sense that we use the term (Friston, 1997; Deco and Jirsa, 2012; Kringelbach et al., 2015; Stratton and Wiles, 2015; Deco and Kringelbach, 2016; Balaguer-Ballester et al., 2018).

The overall dynamic stability of a system may be estimated by calculating a well-defined collective variable or order parameter (Kuramoto, 1984; Shanahan, 2010; Cabral et al., 2011; Wildie and Shanahan, 2012). The Kuramoto order parameter captures the average phase of a group of oscillators to quantify how ‘phase-locked’ they are at a given moment in time. Accordingly, the variation in this order parameter has been proposed as a measure of a system’s metastability and the mean of the phase-locking across time as a measure of the system’s overall synchrony. Metastability is high in a system that visits a range of different states over time whereas both highly ordered and highly disordered states are characterised by low metastability, and high and low phase synchrony, respectively. The concept of phase synchronisation was originally introduced in physics to study the behaviour of weakly coupled oscillators (Rosenblum et al., 1996). The original motivation was to compare the temporal structure of two time series by ignoring information related to amplitude (Varela et al., 2001). Signal processing techniques such as the Hilbert transform make it possible to separate a time series into its amplitude and phase by converting the real signal into its complex analytic version (Boashash, 1992). However, unlike correlation-based sliding-window analysis, which mandates an arbitrary choice of window length, the phase synchronisation approach provides time-resolved functional connectivity at the same resolution as the input narrowband fMRI signal (Glerean et al., 2012). Moreover, unlike correlation, which is a linear measure of association between variables, phase synchronisation is a measure of statistical dependence that is sensitive to both linear and nonlinear relationships (Pereda et al., 2005). Recently, the phase synchronisation approach has successfully identified changes in the time-varying properties of brain connectivity associated with several neural disorders (Hellyer et al., 2015; Córdova-Palomera et al., 2017; Demirtaş et al., 2017; Alderson et al., 2018; Koutsoukos and Angelopoulos, 2018; Lee et al., 2018).

Theoretical accounts stipulate that metastability at rest corresponds to an optimal exploration of the dynamical repertoire inherent in the static structural linkages of the anatomy where the probability of network switching is maximal (Cabral et al., 2011; Ponce-Alvarez et al., 2015; Deco et al., 2017). A critical next step in our understanding is to evaluate not only the degree of metastability arising spontaneously from the brain’s intrinsic network dynamics but also the degree of metastability engendered by the attendant demands of a task (Fingelkurts and Fingelkurts, 2001; Rabinovich et al., 2008). Here, we invoke the theoretical framework of metastable coordination dynamics to explain how resting state networks are dynamically linked into task-dependent neurocognitive networks. Given that patterns of brain activity appear to be more stable during cognitive operations requiring sustained attention (B. Chen et al., 2015; Elton and Gao, 2015; Hutchison and Morton, 2015; Cohen, 2018), we anticipated reduced metastability between task-relevant neural networks as a function of task performance.

In light of the foregoing, we tested the hypothesis that coupling between the brain’s large-scale networks is more metastable at rest than during the execution of an explicit task. We compared the metastability of fMRI BOLD signal in resting and task-evoked functional MRI data in a large cohort of healthy individuals (N=566) from the Human Connectome Project (Van Essen et al., 2013). Changes in metastability were sought among 13 resting state networks encompassing hundreds of regions and every major brain system (Gordon et al., 2016). Finally, a link between the metastability of individual network connections and task performance was sought across several cognitive domains.

Overall, we found that–contrary to expectations–the metastability of couplings between large-scale networks was actively enhanced by task performance, principally in regions known to be devoted to cognitive control. Moreover, the efficiency of the transformation between rest and task-driven states was promoted by a network structure characterised by dynamic flexibility in cognitive control networks and dynamic stability in sensory regions. Crucially, subjects with resting state architectures similar or ‘pre-configured’ to a task-orientated configuration demonstrated superior cognitive ability. Curiously, task-induced increases in metastability did not account for variations in cognitive performance. Rather, cognitive ability was linked to the metastability of the brain’s intrinsic network dynamics. Overall, high intrinsic metastability of cognitive control networks was linked to novel problem solving (or fluid intelligence) but was less relevant in tasks dependent upon previous knowledge and experience (or crystallised intelligence).

## Methods

### Participants

Data were obtained through the Washington University-Minnesota Consortium Human Connectome Project (HCP; Van Essen et al., 2013). Subjects were recruited from Washington University and surrounding area. The present paper used 566 subjects from the 1200 healthy young adult release (aged 22-35; see https://www.humanconnectome.org/data). All participants were screened for a history of neurological and psychiatric conditions and use of psychotropic drugs, as well as for physical conditions or bodily implants. Diagnosis with a mental health disorder and structural abnormalities (as revealed by fMRI) were also exclusion criteria. All participants supplied informed consent in accordance with the HCP research ethics board. The subset of subjects comprising monozygotic and dizygotic twin pairs were excluded from the present study. All subjects attained at a minimum a high school degree.

### MRI parameters

In all parts of the HCP, participants were scanned on the same equipment using the same protocol (Smith et al., 2013). Whole-brain echoplanar scans were acquired with a 32 channel head coil on a modified 3T Siemens Skyra with TR = 720 ms, TE = 33.1 ms, flip angle = 52°, BW = 2290 Hz/Px, in-plane FOV = 208×180 mm, 72 slices, 2.0 mm isotropic voxels, with a multi-band acceleration factor of 8. Rest (eyes open with fixation) and task-based fMRI data were collected over two sessions. Each session consisted of two rest imaging sessions of approximately 15 minutes each, followed by task-based acquisitions of varying length. The present study used only the first resting state run. Except for the run duration, task-based data were acquired using the same EPI pulse sequence parameters as rest. Seven tasks totalling one hour were acquired. Three tasks were collected in one session and four tasks another. The seven tasks (and their run times in mins) were as follows: emotion perception (4:32), relational reasoning (5:52), language processing (7:54), working memory (10:02), gambling/reward learning (6:24), social cognition (theory of mind; 6:54), and motor responses (7:08; Barch et al., 2013). High-resolution 3D T1-weighted structural images were also acquired with the following parameters: TR = 2400 ms, TE = 2.14 ms, TI = 1000 ms, flip angle = 8°, BW = 210 Hz/Px, FOV = 224×224, and 0.7 mm isotropic voxels.

### Task protocols

Task-evoked fMRI data were downloaded to examine the changes in metastable interactions between large-scale cortical networks during attentionally demanding cognition. In total there were seven in-scanner tasks designed to engage a variety of cortical and subcortical networks related to emotion perception, relational reasoning, language processing, working memory, gambling/reward learning, social cognition (theory of mind), and motor responses. These included:

**1. Emotion perception**: participants were presented with blocks of trials asking them to decide which of two faces presented on the bottom of the screen matched the face at the top of the screen, or which of two shapes presented at the bottom of the screen matched the shape at the top of the screen. The faces had either an angry or fearful expression (Hariri et al., 2002).
**2. Relational reasoning**: participants were presented with 2 pairs of objects, with one pair at the top of the screen and the other pair at the bottom of the screen. Subjects were first asked to decide if the top pair of objects differed in shape or differed in texture and then to decide whether the bottom pair of objects also differed along the same dimension (Smith et al., 2007).
**3. Language processing**: the task comprised a story and math component. The story blocks presented participants with brief auditory stories (5-9 sentences) adapted from Aesop’s fables, followed by a 2-alternative forced-choice question that asked participants about the topic of the story. The math task also presented trials auditorily and required subjects to complete addition and subtraction problems (Binder et al., 2011).
**4. 2-back working memory**: task participants were presented with blocks of trials that consisted of pictures of places, tools, faces and body parts (non-mutilated, non-nude). The task consisted of indicating when the current stimulus matched the one from 2 steps earlier.
**5. Gambling/reward learning**: participants were asked to guess the number on a mystery card in order to win or lose money. Participants were told that potential card numbers ranged from 1-9 and that the mystery card number was more than or less than 5 (Delgado et al., 2000).
**6. Social cognition (theory of mind)**: participants were presented with short video clips (20 seconds) of objects (squares, circles, triangles) that either interacted in some way, or moved randomly on the screen. After each video clip, participants were asked to judge whether a mental interaction had occurred; did the shapes appear to take into account each other’s thoughts and feelings? (Wheatley et al., 2007; Castelli et al., 2013).
**7. Motor responses**: participants were presented with visual cues that asked them to either tap their left or right fingers, squeeze their left or right toes, or move their tongue (Buckner et al., 2011; Yeo et al., 2011).

Full timing and trial structure for the seven tasks are provided as supplementary information.

### Task fMRI behavioural data

Task performance was evaluated using behavioural accuracy and reaction time data. Only those tasks which showed normally distributed behavioural accuracy scores were utilised for subsequent analysis. As confirmed by a Kolmogorov-Smirnov test (p<0.05) three of the seven tasks satisfied this criteria including relational reasoning (M=0.76, SD=0.12), language processing (M=0.88, SD=0.71), and working memory (M=0.83, SD=0.10). Of the other four tasks, the gambling task accuracies were no better than chance (participants were asked to guess if a mystery card was higher or lower than five). The emotion (M=0.97, SD=0.03) and social (M=0.96, SD=0.12) task accuracies were perfect or near perfect for most subjects and hence showed a strong ceiling effect. Finally, the motor task accuracy scores were not recorded (subjects were asked to move tongue, hands, or feet). All seven tasks showed normally distributed reaction time data, as confirmed by a one-sample Kolmogorov-Smirnov test (p<0.05).

### Cognitive measures

Cognitive performance was also evaluated using test scores obtained outside the scanner. These included two complementary factors of general intelligence: fluid and crystallised intelligence, the former linked to novel problem solving and the latter to previously acquired knowledge and experience (Jensen and Cattell, 2006). Executive function/inhibitory control was also investigated due to its strong association with tonic (Sadaghiani and D’Esposito, 2015) and phasic (Cole et al., 2012, 2013) aspects of attention.

The HCP provides a comprehensive and well-validated battery of cognitive measures based on tools and methods developed by the NIH Toolbox for Assessment of Neurological and Behavioural Function (Gershon et al., 2013). Relevant cognitive measures were downloaded from the Connectome Database (https://db.humanconnectome.org; Hodge et al., 2015). These included fluid intelligence (Penn Progressive Matrices; PMAT), crystallised intelligence (NIH Toolbox Picture Vocabulary Test and NIH Toolbox Oral Reading Recognition Test), and executive function/inhibitory control (NIH Toolbox Flanker Inhibitory Control and Attention Test). All three cognitive measures were consistent with a normal distribution as confirmed by a one-sample Kolmogorov-Smirnov test (p<0.05).

### fMRI pre-processing

All pre-processing was conducted using custom scripts developed in MATLAB 2017a (The MathWorks, Inc., Natick, Massachusetts, United States). Motion between successive frames was estimated using framewise displacement (FD) and root mean square change in BOLD signal (DVARS; Power et al., 2012, 2014, 2015; Burgess et al., 2016). FD was calculated from the derivatives of the six rigid-body realignment parameters estimated during standard volume realignment. If more than 20% of a subject’s resting state frames exceeded FD > 0.5 mm or DVARS > 5%, they were excluded from further analysis. Based on this criteria, 566 out of 890 subjects were retained for further analysis.

We used a minimally pre-processed version of the data that included spatial normalization to a standard template, motion correction, slice timing correction, intensity normalization, and surface and parcel constrained smoothing of 2 mm full width at half maximum (Glasser et al., 2013). The data corresponded to the standard ‘grayordinate’ space consisting of left and right cortical surface meshes and a set of subcortical volume parcels which have greater spatial correspondence across subjects than volumetrically aligned data (Glasser et al., 2016). To facilitate comparison between rest and task-based conditions both sets of data were identically processed. T1 weighted images aligned with the standard functional image were segmented into 3D volume masks with Freesurfer (Fischl et al., 2002, 2004). Average signals were extracted from the voxels corresponding to the ventricles and white matter anatomy. Variables of no interest were removed from the time series by linear regression. These included six linear head motion parameters, mean ventricle and white matter signals, and corresponding derivatives.

Since each task comprised two runs (one from each session) both runs were concatenated into a single time series. The individual signals were demeaned and normalised by z-scoring the data. To pre-empt the possibility that variation in synchrony (our definition of metastability) was being driven by alternating blocks of task and fixation, task blocks were concatenated. Since artificially concatenating a series of disjoint task blocks resulted in a discontiguous time series, the analysis was also performed with cue and fixation blocks included. Overall, retaining cue and fixation blocks did not alter the pattern of metastability between large-scale networks (only the statistical significance). The present analysis pertains to the case where cue and fixation blocks are removed. To ensure that any observed differences were due to dynamics rather than bias associated with signal length, the same number of contiguous frames from task and rest were utilised; the resting state scan was truncated to match the length of the task run (after cue and fixation blocks were removed).

To obtain meaningful signal phases and avoid introducing artefactual correlations, the empirical BOLD signal was bandpassed filtered (Glerean et al., 2012). Since low frequency components of the fMRI signal (0–0.15 Hz) are attributable to task-related activity whereas functional associations between high frequency components (0.2–0.4 Hz) are not (Sun et al., 2004), a temporal bandpass filter (0.06–0.125 Hz) was applied to the data (Shine et al., 2016a). The frequency range 0.06–0.125 Hz is thought to be especially sensitive to dynamic changes in task-related functional brain architecture (Bassett et al., 2011, 2013, 2015; Glerean et al., 2012).

### Brain parcellation

Mean time series were extracted from regions of interest defined by the Gordon atlas (Gordon et al., 2016). The separation of regions into functionally discrete time courses is especially suitable for interrogating dynamic fluctuations in synchrony between large-scale networks. The Gordon atlas assigns regions to one of 12 large-scale networks corresponding to abrupt transitions in resting state functional connectivity. These include dorsal attention, ventral attention, fronto-parietal, cingulo-opercular, salience, default mode, medial parietal, parietal-occipital, visual, motor mouth, motor hand, and auditory networks. Regions outside these domains are labelled as ‘none’. The atlas was downloaded from the Brain Analysis Library of Spatial Maps and Atlases database (https://balsa.wustl.edu; Van Essen et al., 2017). Whole-brain coverage consisted of 333 cortical regions (161 and 162 regions from left and right hemispheres respectively), and one subcortical volume corresponding to the thalamus. The thalamus, which exhibits domain-general engagement across multiple cognitive functions, also plays a critical role in integrating information across functional brain networks (Hwang et al., 2017).

### Calculating resting state network metastability

The first step in quantifying phase synchronisation of two or more time series is determining their instantaneous phases. The most common method is based on the analytic signal approach (Gabor, 1946; Panter, 1965). The advantage of the analytic signal is that by ignoring information related to amplitude additional properties of the time series become accessible. From a continuous signal *x* (*t*) the analytic signal *x_a_*(*t*) is defined as,

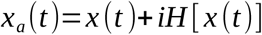

where *H* is the Hilbert transform and 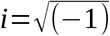. If Bedrosian’s theorem (Bedrosian, 2008) is respected then the analytic signal of a time series can be rewritten as,

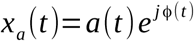

where *a* (*t*) is the instantaneous envelope and ϕ(*t*) the instantaneous phase. The Bedrosian theorem makes a clear prediction–the narrower the bandwidth of the signal of interest, the better the Hilbert transform is able to generate an analytic signal with meaningful envelope and phase. For this reason, bandpass filtering of empirical BOLD signal is essential prior to performing the transform. In accordance with the foregoing, the 334 narrowband mean BOLD time series were transformed into complex phase representation via a Hilbert transform. The first and last ten time points were removed to minimise border effects inherent to the transform (Ponce-Alvarez et al., 2015; Córdova-Palomera et al., 2017).

The ‘instantaneous’ collective behaviour of a group of phase oscillators may be described in terms of their mean phase coherence or synchrony. A measure of phase coherence–the Kuramoto order parameter (Strogatz, 2000; Acebrón et al., 2005) was estimated for: (1) the set of regions comprising a single resting state network; and (2) to evaluate interactions, the set of regions comprising two resting state networks as:

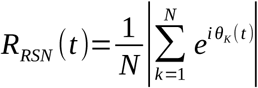

where *k* ={1,... *, N* } is region number and *θ _K_* (*t*) is the instantaneous phase of oscillator *k* at time *t*. Under complete independence, all phases are uniformly distributed and *R_RSN_* approaches zero. Conversely, if all phases are equivalent, *R_RSN_* approaches one and full phase synchronisation. The maintenance of a particular communication channel through coherence implies a persistent phase relationship. The number or repertoire of such channels therefore corresponds to the variability of these phase relationships. Accordingly, metastability is defined as the standard deviation of *R_RSN_* and synchrony as the mean of *R_RSN_* (Shanahan, 2010; Cabral et al., 2011; Deco and Kringelbach, 2016). Global metastability was estimated by considering the interactions of all resting state networks simultaneously (all 334 cortical and subcortical signals).

### Assessing changes in empirical resting state network metastability during tasks

Given that, (1) the current formulation of metastability permits calculation between a group of regions; and (2) interconnected subnetworks convey more behaviourally relevant information than functional connections observed between pairs of regions observed in isolation, we advocate for a method that exploits the clustering structure of connectivity alterations between functionally related networks. For this reason, we applied the network-based statistic (NBS) to estimates of empirical metastability obtained from fMRI data at the network rather than regional level (see also, Alderson et al., 2018). For each subject, we estimated an ‘interaction matrix’ reflecting the metastable interactions of the 13 resting state networks (and thalamus) defined by the Gordon atlas. The same procedure was applied to compute an equivalent interaction matrix based on synchrony.

The NBS is a non-parametric statistical test designed to deal with the multiple comparisons problem by identifying the largest connected sub-component (either increases or decreases) in topological space while controlling the family wise error rate (FWER). To date, several studies have used the method to identify pairwise regional connections that are associated with either an experimental effect or between-group difference in functional connectivity (Zalesky et al., 2010). Here, we use the NBS to identify topological clusters of altered metastability (or synchrony) between empirical resting state networks under different conditions of task relative to rest.

Mass univariate testing was performed at every connection in the graph to provide a single test statistic that captured the evidence in favour of the null hypothesis: that there was no statistically significant difference in the means of resting state and task-based metastability. The test statistic was subsequently thresholded at an arbitrary value with the set of supra-threshold connections forming a candidate set of connections for which the null hypothesis was tested. Topological clusters were identified between the set of supra-threshold connections for which a single connected path existed. The null hypothesis, therefore, was accepted or rejected at the level of the entire connected graph rather than at the level of an individual network connection. The above steps were repeated in order to construct an empirical null distribution of the largest connected component sizes. Finally, FWE-corrected p-values, corresponding to the proportion of permutations for which the largest component was of the same size or greater, were computed for each component using permutation testing.

### Classification of task and rest data

The interaction matrices corresponding to the seven different tasks (plus rest) were classified using a modified convolutional neural network (CNN) architecture (Fig. 1). BrainNetCNN is the first deep learning framework designed specifically to leverage the topological relationships between nodes in brain network data, outperforming a fully connected neural network with the same number of parameters (Kawahara et al., 2017). The architecture of BrainNetCNN is motivated by the understanding that local neighbourhoods in connectome data are different from those found in traditional datasets informed by images. Patterns are not shift-invariant (as is a face in a photograph) and the features captured by the local neighbourhood (e.g. a 3×3 convolutional filter) are not necessarily interpretable when the ordering of nodes is arbitrary.

**Figure 1:**
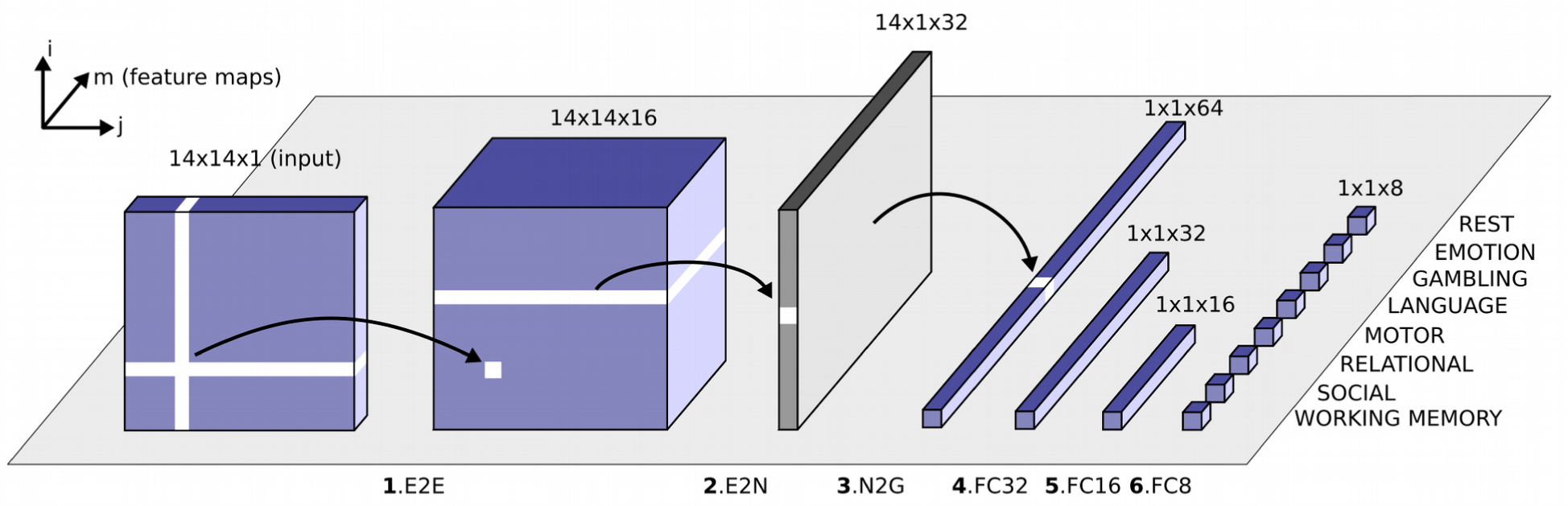
Schematic overview of the modified convolutional neural network architecture used to classify the eight different network configurations (seven tasks and one resting state; Kawahara et al., 2017). Each block represents the input/output of a numbered filter layer. The third dimension (m) represents the result of convolving the input with m different filters (feature maps). First, an interaction matrix composed of the interactions of 14 networks (based on synchrony or metastability) is entered as input. This is convolved with an edge-to-edge (1. E2E) filter which weights the edges associated with adjacent brain networks in topological space. The output from this layer is then convolved with an edge-to-node (2. E2N) filter which assigns each network a weighted sum of its edges. Next, a node-to-graph (3. N2G) layer outputs a single response based on all the weighted nodes. Finally, the number of features is reduced to eight output classifications through a series of fully connected (4/5/6. FC) layers.

To reduce the number of parameters we included only a single edge-to-edge layer (Meszlényi et al., 2017). The input to the CNN is the set of 14×14 interaction matrices that capture the metastability/synchrony between the 13 resting state networks (plus the thalamus) defined by the Gordon atlas. The network classifies the data into one of the seven tasks or the subject’s resting state (random classification accuracy is 12.5%). The model was evaluated using k-fold cross validation where *k* =5. This value of *k* has been shown to yield test error rate estimates that suffer neither from excessively high bias nor from very high variance (Kuhn and Johnson, 2013).

The original dataset was partitioned randomly into training (60%), validation (20%), and testing sets (20%). That is, 340 subjects were assigned for training the model, 113 subjects were assigned for tuning the model’s hyperparameters, and a further 113 were withheld for validating the performance of the trained model. In the case of metastability, each of the 566 subjects was associated with 8 interaction matrices (7 task-based interaction matrices and one resting state interaction matrix). The same was true in the case of synchrony. Performance was evaluated using classification accuracy i.e. the proportion of correctly identified instances. The above procedure was repeated twice, once for the interaction matrices capturing metastability and again using the interaction matrices based on synchrony.

The CNN was implemented in Python using the Pytorch framework (Paszke et al., 2017). Rectified linear units (RELUs; Nair and Hinton, 2010) were used as activation functions between layers and the probability of each class was calculated at the output layer using the soft max function (Bridle, 1990). The network was trained using the Adam optimiser (Kingma and Ba, 2015) with mini-batch size of 128, a learning rate of 0.001, and momentum of 0.9. Drop-out regularisation of 0.6 was applied between layers to prevent over-fitting (Wager et al., 2013; Srivastava et al., 2014). The model minimised a cost function associated with the cross-entropy loss. Hyperparameters used in the optimisation stage included momentum and drop-out regularisation.

### Defining update/reconfiguration efficiency

The ability to switch from a resting state network architecture into a task-based configuration was designated as update efficiency (Schultz and Cole, 2016). Highly similar rest and task-based network configurations are commensurate with high update efficiency, as few changes are required to transition between the two whilst highly dissimilar resting state and task-based architectures are linked to low update efficiency, reflecting the many changes that are required to make the switch. Update efficiencies were calculated for all 566 subjects by vectorising the upper triangular half and diagonal of the rest and task-general interaction matrices and calculating their Pearson’s correlation coefficient.

## Results

### Higher global metastability during task than rest

The metastability of fMRI BOLD signal was examined during the resting state and during the execution of several cognitively demanding tasks (Fig. 2). One-way ANOVA identified a statistically significant difference between groups (F(7,4520) = 37.32, p < 0.01). Subsequent Tukey post hoc test revealed significantly higher global metastability during the seven tasks as compared the resting state (M = 0.112, SD = 0.019). These included emotion perception (M = 0.132, SD = 0.031), relational reasoning (M = 0.133, SD = 0.030), language processing (M = 0.134, SD = 0.027), working memory (M =0.135, SD = 0.027), gambling/reward learning (M = 0.136, SD = 0.029), social cognition (M = 0.141, SD = 0.029), and motor responses (M = 0.143, SD = 0.030).

**Figure 2:**
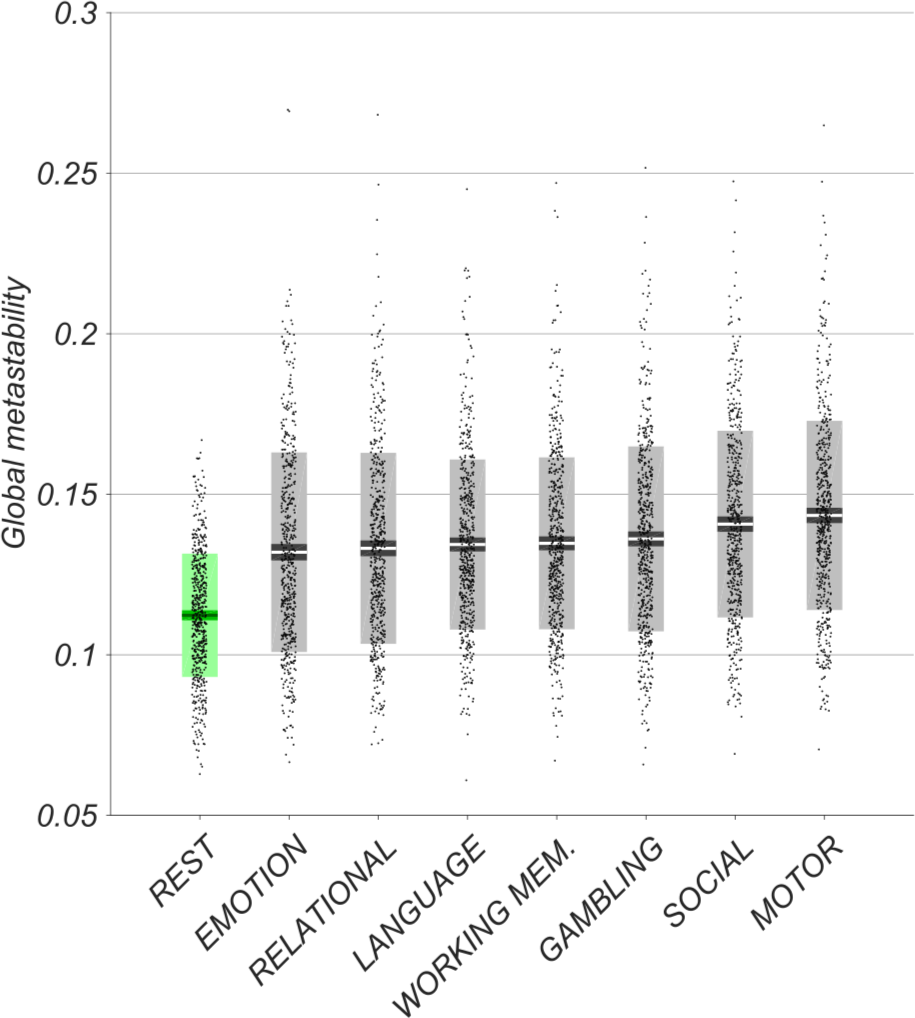
Empirical global metastability of fMRI BOLD signal in the resting state (in green) and during several cognitively demanding tasks (in grey). Bars display mean, 95% CI, and one standard deviation with individual subjects indicated. Tasks arranged in ascending order of mean metastability. One-way ANOVA revealed significantly higher metastability during task execution relative to resting state (p < 0.01).

### Task related increases in metastability between resting state networks

The NBS was subsequently used to identify changes in the metastability of fMRI BOLD signal of individual network connections (task versus rest). In total, 566 resting state interaction matrices were compared to 566 task-based interaction matrices within each of the seven behavioural domains. The null hypothesis, that there was no difference in metastability between rest and task, could then be rejected at the level of individual network connections.

Consistent with the role of resting state networks in mediating behaviour (Sadaghiani, 2010; Sadaghiani and Kleinschmidt, 2013), the NBS identified statistically significant (p < 0.01; corrected) increases in metastability between several large-scale networks during task engagement relative to the more unconstrained resting state. Figure 3 shows the largest connected sub-graph of increased metastability detected by the NBS at a fixed threshold for all seven tasks where each node is scaled to reflect its relative importance within the sub-graph (the sum of its effect sizes). Cortical regions associated with top-down attentional/cognitive control (including fronto-parietal and dorsal attention networks) were the most active across the seven tasks along with thalamo-cortical interactions linked to memory, learning, and flexible adaptation (Alcaraz et al., 2018; Wolff and Vann, 2018).

**Figure 3:**
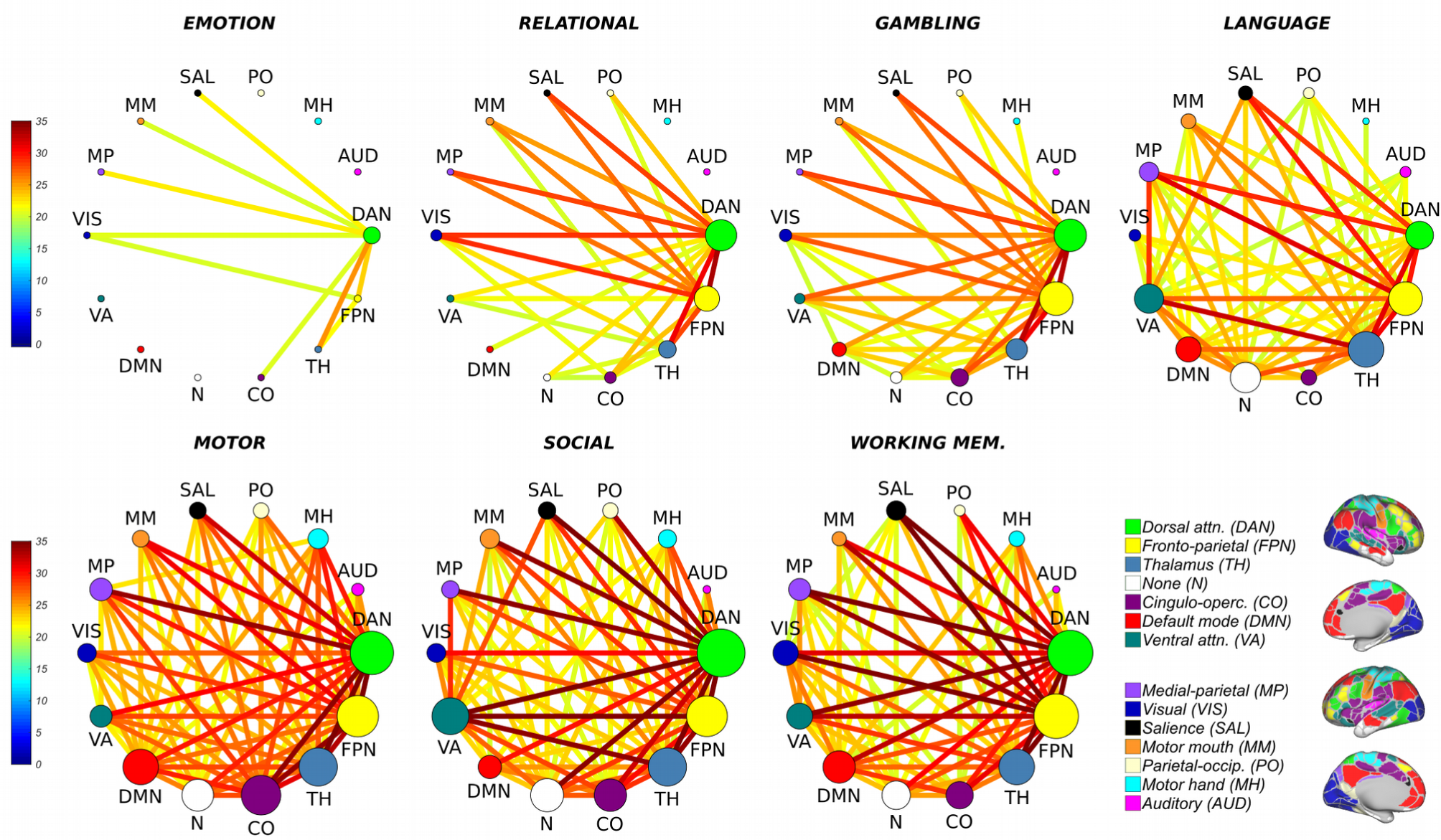
Statistically significant (p < 0.01; corrected) increases in BOLD signal metastability between empirical resting state networks during task as compared to resting state. Circular graphs show largest connected sub-graph of increased metastability identified by the network-based statistic at a fixed threshold (16). Nodes are scaled to reflect the relative importance of their interactions (the sum of their effect sizes). Overall, the connectivity of the dorsal attention (green) and fronto-parietal networks (yellow) were the most metastable.

### Increases in metastability were more widespread than equivalent increases in synchrony

Edges associated with each sub-graph (metastability and synchrony) were summed to reveal the total number of network connections recruited by each task (Fig. 4). Increases in metastability spanned a greater number of cognitive subsystems than equivalent increases in synchrony (in all but emotion and motor tasks). In both cases, the NBS received an identical threshold.

**Figure 4:**
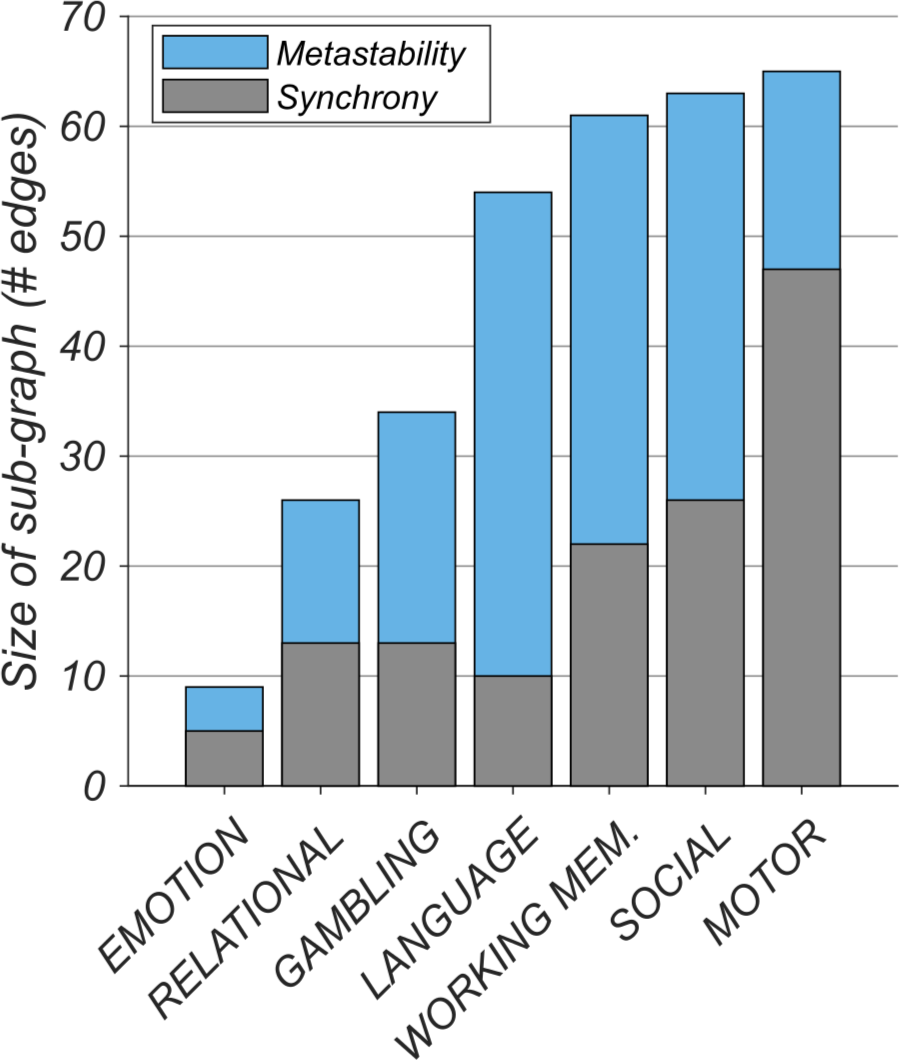
Increases in metastability (blue) were associated with a greater number of network connections than equivalent increases in synchrony (grey). Figure shows size of sub-graph identified by the network-based statistic (see Fig. 3) rank ordered by metastability.

### Each task is characterised by a small number of task-evoked changes in synchrony

The highly correlated properties of metastability and synchrony (see Fig. S1 in the supplementary information) were disassociated using a deep learning framework. Accordingly, task and rest network states captured by the 14×14 interaction matrices of metastability and synchrony were provided as input to a CNN for classification. The network correctly identified the 7 different tasks (plus rest) based on synchrony with high accuracy (76% average; chance level 12.5%; Fig. 5.A) but performed less well when trained on metastability (46% average; Fig. 5.B). The high sensitivity (true positive rate) and specificity (true negative rate) exhibited by the classifier when trained on synchronous interactions between networks suggested that each behavioural domain was characterised by a small number of unique task-evoked network changes. This was confirmed by masking out inputs (interactions between networks) relevant for correct classification (Fig. 6) and re-evaluating the pre-trained classifier. Overall, occluded inputs were associated with exceptionally poor classification accuracy (Fig. 5.C).

**Figure 5:**
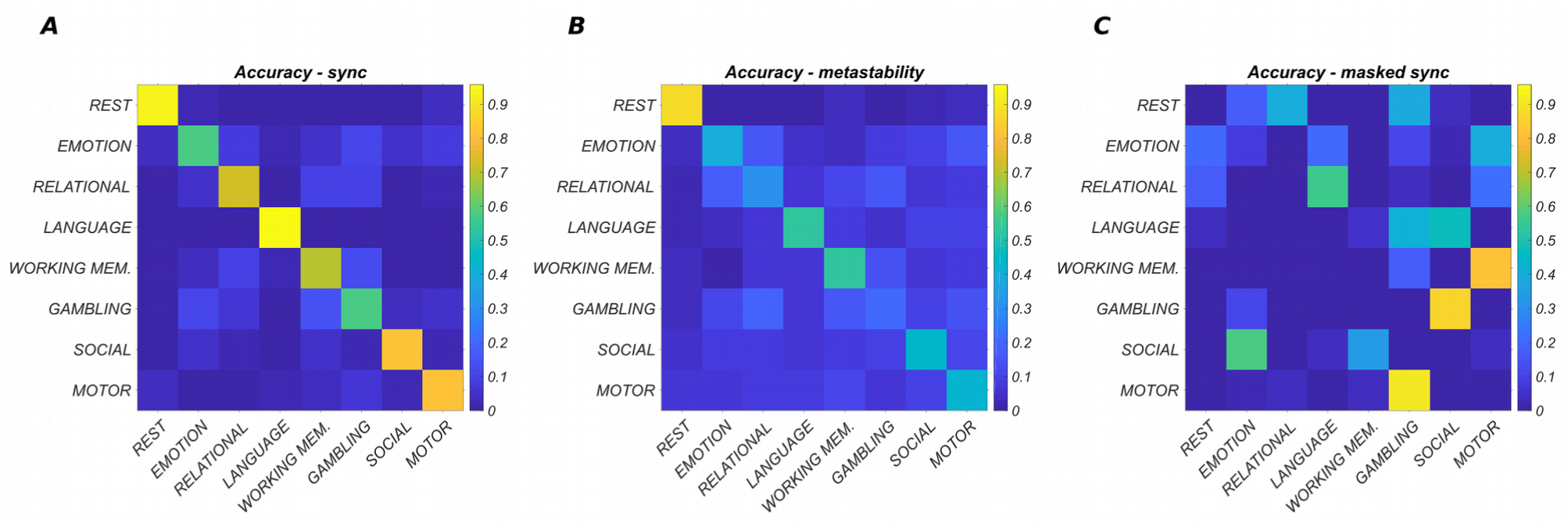
Convolutional neural network performance in terms of classification accuracy where each row represents the true class and each column represents the classification made by the neural network. Diagonal elements report the percentage of instances correctly classified. Off-diagonal elements report the percentage of instances that are incorrectly classified. Inputs were classified as belonging to one of eight different network states (seven tasks plus one resting state condition) where each row/column corresponded to the interaction of one network with 13 others (in terms of either synchrony or metastability). **A**, Classification accuracy in terms of the synchrony between networks (average accuracy = 76%; chance level 12.5%). **B**, Classification accuracy in terms of the metastability between networks (average accuracy = 46%). **C**, Classification accuracy in terms of occluded network synchrony (average accuracy = 2%). Here, classification accuracy was reduced by masking out (setting to zero) a small subset of network interactions (see Fig. 6).

**Figure 6:**
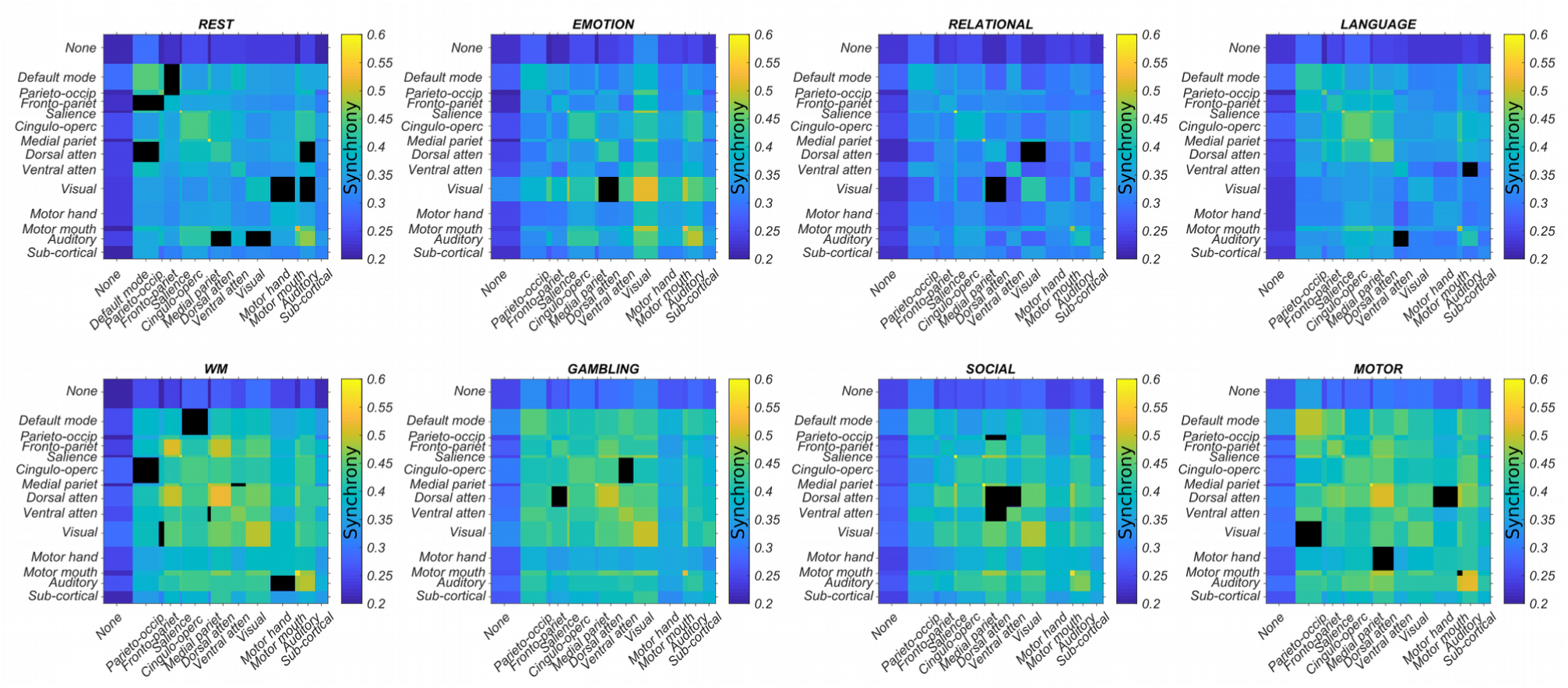
Each network state (one rest and seven tasks) is defined by a small number of task-evoked changes in synchrony between resting state networks. Here, network connectivity important for correct classification in more than 90% of individuals (as determined by guided backpropagation) is masked out (black). Such ‘occluded’ inputs are associated with exceptionally poor classification accuracy (Fig. 5C). The width of each column/row has been scaled to reflect the relative number of regions in each network.

In detail, guided backpropagation was used to identify the most important inputs for correctly classifying each task. Guided backpropagation provides a set of gradients relating input to output. High gradients at the input level have a large effect on the output and are therefore important for classification. Recall, that each row/column of the 14×14 input represented the interaction of a single network (and thalamus) with 14 others. Thus, guided backpropagation produced a 14×14 matrix of gradients. A consensus across all subjects for a particular task was obtained by setting each subject’s top 10% of gradients (the most positive gradients) to one and the remaining entries to zero, summing the matrices, and dividing by the total number of subjects. Entries important for correct classification in more than 90% of individuals were set to zero in the input (the interaction matrix). The performance of the pre-trained classifier was then re-evaluated based on the occluded input data. In a separate experiment, retraining on the occluded inputs also produced extremely poor classification accuracy (not shown).

### Different behaviours recruit a similar set of metastable connections

So far, we have demonstrated that increases in metastability can be distinguished from increases in synchrony in two ways: (1) their overall network size, that is, increases in metastability encompass a wider network of cognitive systems than those based on synchrony; and (2) their discriminatory utility, that is, tasks can be identified with high accuracy based on a small subset of network changes in synchrony (but much less so in terms of metastability). Taking these findings as a whole, we hypothesised that commonalities between tasks may be centred around a metastable core of task-general network interactions.

To quantify the degree to which the seven task-based configurations shared features in common, we used a dimension-reduction tool–principal component analysis (PCA)–to reduce a larger set of variables (the seven task-based interaction matrices) to a smaller set (a single task-general network architecture) retaining most of the information. Accordingly, entering the seven task-based interaction matrices based on metastability into a PCA yielded a single task-general architecture for each subject. On average, the first principal component accounted for 78% of the variance between the seven tasks. The loadings were positive and distributed equally between the seven tasks suggesting that each task-based configuration contributed equally to the task-general structure. These included emotion perception=0.37, relational reasoning=0.37, language processing=0.36, working memory=0.36, gambling/reward learning=0.36, social cognition=0.37, and motor responses=0.38. This result likely speaks to the high similarity between task-based configurations engaged by different behaviours (see Fig. S2 in the supplementary information). An exemplar task-general architecture was subsequently derived through simple averaging across subjects.

### Task general architecture is composed of high and low metastability subnetworks

We explicated this structure by performing another PCA analysis on the interaction matrices obtained by subtracting task from rest. In doing so, the task general architecture was decomposed into both high (Fig. 7; top) and low (Fig. 7; middle) metastability subnetworks. High metastability (Fig. 7; red) was found in networks associated with cognitive control including dorsal attention (selective attention in external visuospatial domains) and fronto-parietal networks (adaptive task control). In contrast, low metastability (Fig. 7; blue) was linked to unimodal (or modality specific) sensory processing architecture including motor, auditory, and visual networks. From one perspective, this result is quite surprising. One would expect motor cortex to shift from rest to task due to motor demands on task but primary sensory regions–including motor cortex–appear to favour dynamic stability. Tertiary thalamo-cortical contributions were also apparent (memory, category learning, and adaptive flexibility).

**Figure 7:**
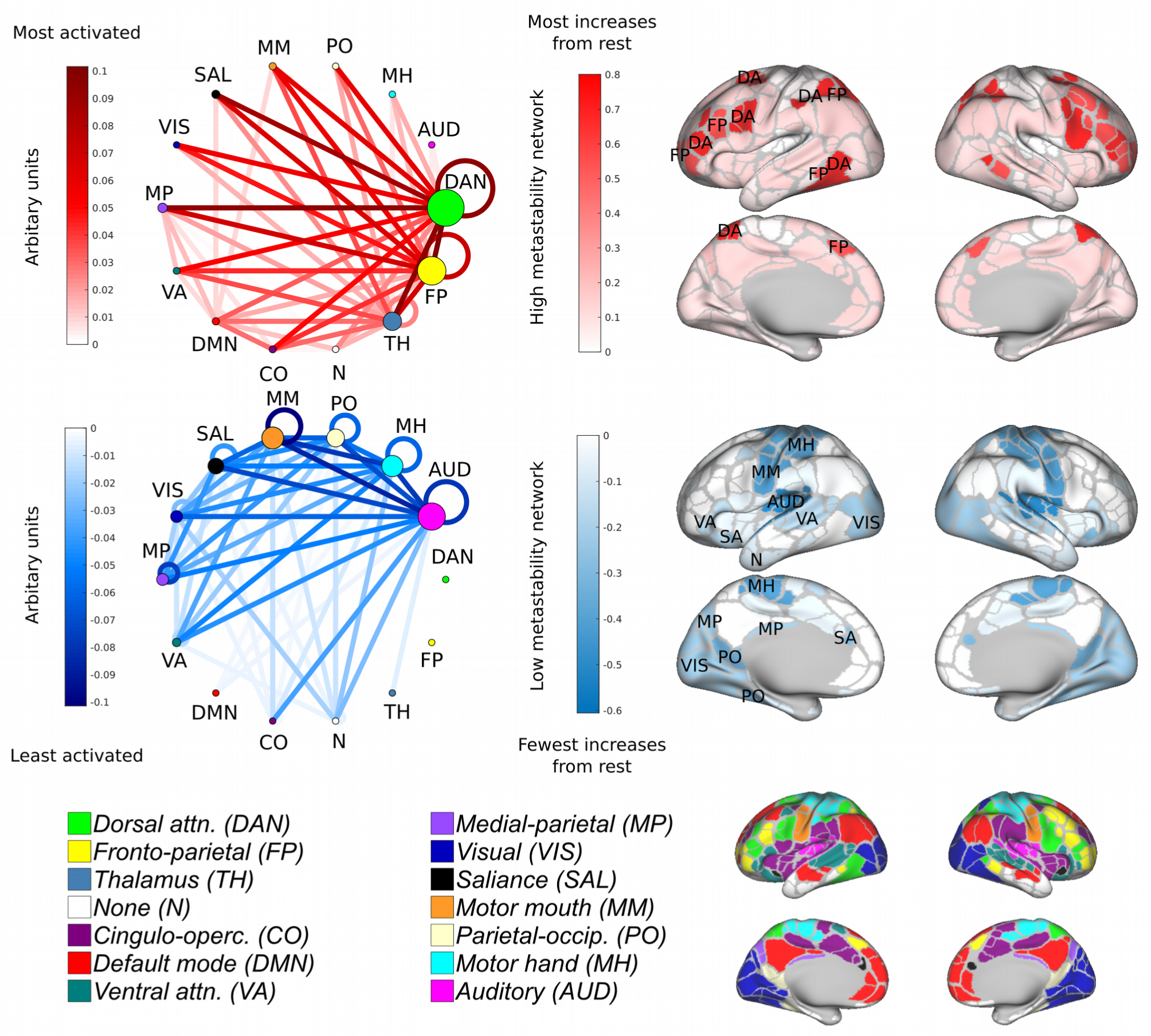
Principle component analysis reveals a task-general network architecture. Task-based reasoning was principally characterised by high metastability in regions associated with cognitive control (top; red) and low metastability in regions associated with sensory processing (blue; middle). On average the 1st principal component accounted for 78% of the variance. Loadings were distributed equally between the seven tasks. Regions are colour coded by the sum of their ingoing/outgoing connectivity. Nodes are colour coded according to the Gordon atlas (bottom). Node diameter is proportional to the sum of ingoing/outgoing connectivity. Recurrent connections correspond to activity within a network.

### High metastability of cognitive control systems at rest is predictive of task performance

We next examined the metastable interactions of large-scale networks during task engagement for evidence that they informed behaviour. Behavioural accuracy scores for each subject were entered into a linear regression analysis as the dependent variable with one of 196 (14×14) task-based network connections (estimated in terms of metastability) as predictors. Overall, metastable interactions between networks during task did not explain the variance in cognitive ability. Resting state metastability was then entered as an additional independent factor. Across the three in-scanner tasks, several network connections demonstrated a significant positive association between intrinsic metastability and cognitive performance (Fig. 8 bottom; p < 0.01; FDR corrected for multiple comparisons). Three additional cognitive measures acquired outside the scanner were also analysed. These included fluid intelligence, crystallised intelligence, and executive function. Across the three measures, several network connections demonstrated a significant positive association between intrinsic metastability and cognitive performance (Fig. 8 top; p < 0.01; FDR corrected for multiple comparisons). The addition of reaction time data across the six tasks did not increase the percentage of explained variance in cognitive performance. In a separate linear regression analysis, no statistically significant associations between behaviour/cognition and network synchrony were identified (p < 0.01; FDR corrected for multiple comparisons).Interactions between networks are presented as circular graphs where each edge represents a significant positive correlation between network metastability and cognition (Fig. 8). In the main, the metastability of large-scale networks related to cognitive control was strongly related to task performance. These included dorsal attention, cingulo-opercular and fronto-parietal networks, and to a lesser extent the salience and ventral attention networks. In general, the metastability of networks related to primary sensory processing (including motor, auditory, and visual networks) was less relevant to cognitive ability. These results are broadly consistent with the task-general network architecture previously discussed.

**Figure 8:**
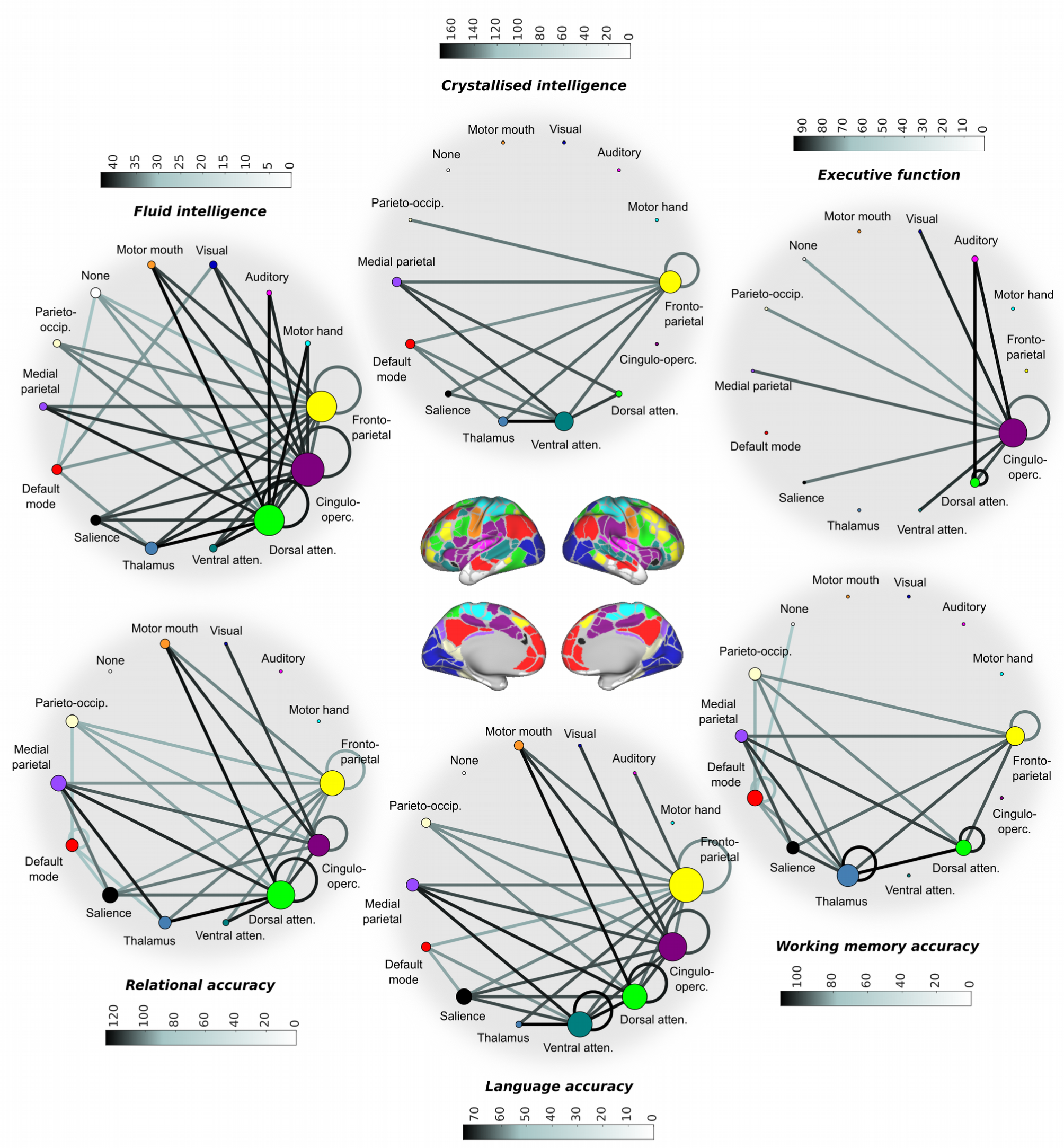
Resting state metastability of cognitive control networks is predictive of task performance. Each edge represents a statistically significant positive correlation between the intrinsic metastability of a connection and cognition/behaviour (p < 0.01; FDR corrected) shaded to reflect standardised effect sizes (Pearson’s r or correlation coefficients). Nodal diameter is scaled to reflect the sum of their ingoing/outgoing connectivity. Cognitive measures were obtained outside the scanner (top); measures of behavioural accuracy were acquired inside the scanner (bottom). Metastability in the connectivity of cognitive control networks was linked to task performance including the fronto-parietal (adaptive task control), cingulo-opercular (sustained tonic attention), and dorsal attention networks (attending to visuospatial stimuli). Note the markedly different profiles presented by fluid and crystallised intelligence.

Concerning specific tasks, the metastability of connections associated with cognitive control networks was strongly related to fluid intelligence. These included dorsal attention, cingulo-opercular, and fronto-parietal networks. In contrast, the metastability of these networks was less important in the execution of crystallised intelligence, which presented a more circumscribed network of significant positive correlations. Finally, executive function/inhibitory control was most strongly associated with the metastability of cingulo-opercular network connections.

Compared to the other in-scanner tasks (relational reasoning and language processing) the working memory task displayed a strong association between metastability and behavioural accuracy in the thalamus. Language processing was distinguished from relational reasoning and working memory by virtue of the association between metastability and cognition in the connectivity of the ventral attention network. Overall, all six behavioural domains, including relational reasoning, language processing, working memory, fluid intelligence, crystallised intelligence, and executive function/inhibitory control, demonstrated significant positive correlations between cognitive performance and spontaneous metastability in regions associated with cognitive control (including the dorsal attention, cingulo-opercular, and fronto-parietal networks).

### High metastability within (and between) cognitive control systems at rest promotes efficient switching into task

Even though it is self-evident that for high update efficiency to be achieved, a subject’s resting and task-general architectures must be in agreement, not all of these relationships will necessarily hold statistically at the chosen significance level (p < 0.01; FDR). For this reason, we correlated the metastability of individual network connections with update efficiency and obtained the slope of the regression equation (Fig. 9.A) and its significance (Fig. 9.B).

**Figure 9:**
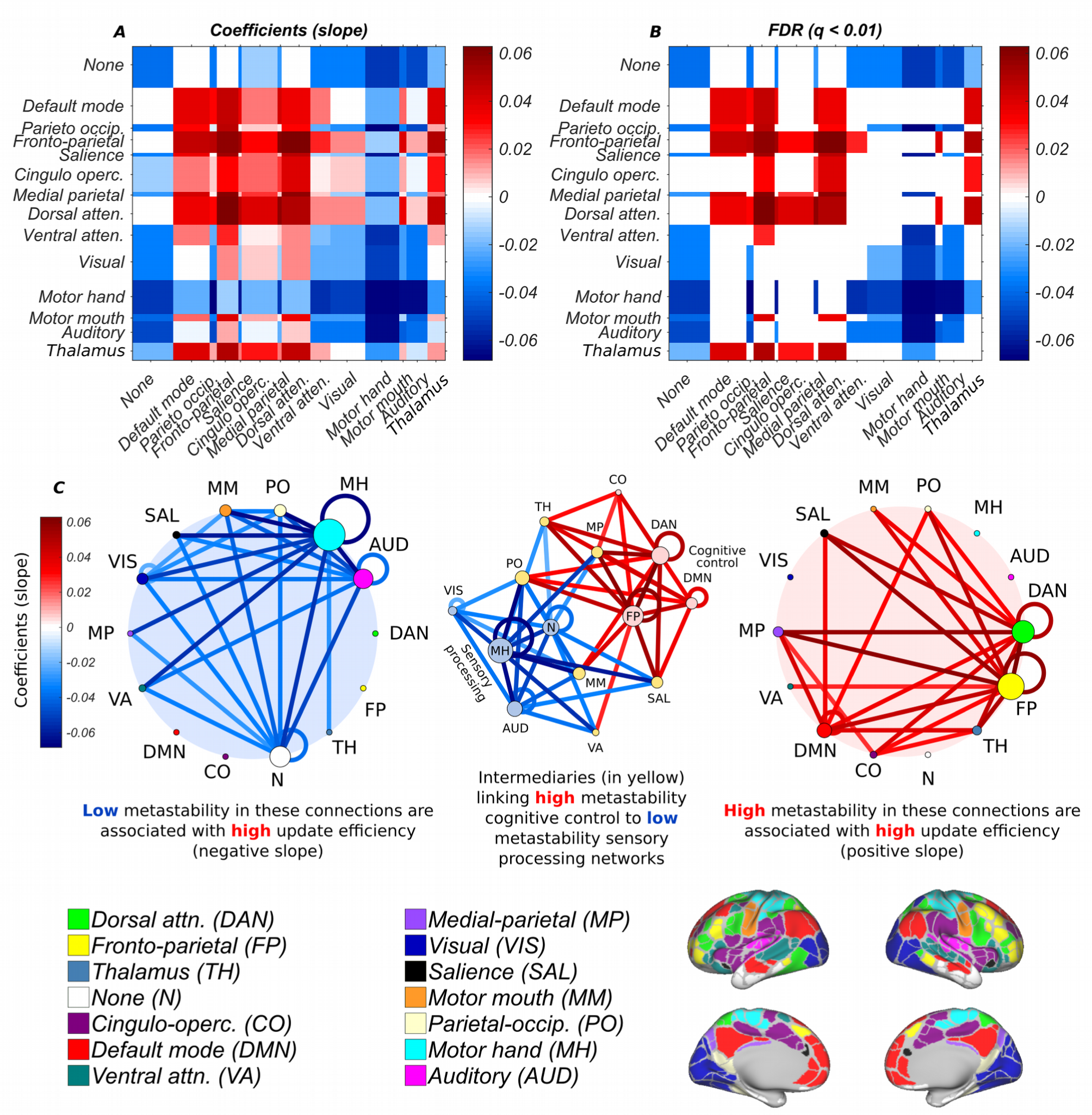
The efficiency of the transformation between resting and task-based network architecture is conditioned on high metastability in the couplings of cognitive control networks and low metastability in the couplings of sensory networks. **A**, Slope (coefficients) of the regression line (negative or positive) relating metastability of network connectivity to update efficiency. **B**, Statistically significant correlations between metastability of network connectivity and update efficiency (p < 0.01; FDR corrected). **C**, Statistically significant correlations between metastability and update efficiency in low (blue; left) and high (red; right) metastability subnetworks (p < 0.01; FDR corrected) where low metastability is associated with unimodal sensory networks (auditory, motor, and visual) and high metastability is related to cognitive control (dorsal attention, fronto-parietal, cingulo-opercular, and default mode networks). Some networks such as the salience, medial parietal, parieto-occipital, and thalamus were sites of convergence for both high and low metastability connections (yellow; centre). Nodal diameter has bee scaled to reflect the sum of their ingoing/outgoing edges.

A significant positive relationship between update efficiency and metastability was found in the fronto-parietal, dorsal attention, cingulo-opercular, and default mode networks (Fig. 9.C; right; red), and a significant negative relationship in the motor, auditory, and visual networks (Fig. 9.C; left; blue; p < 0.01; FDR corrected for multiple comparisons). As expected, these results accord with the structure of the task-general configuration (Fig. 7). Overall, high update efficiency was characterised by dynamic flexibility in the connectivity of networks implicated in cognitive control and dynamic stability in primary sensory network. Some networks such as the salience, ventral attention, medial parietal, parieto-occipital, and thalamus were sites of convergence for both high and low metastability connections (Fig. 9.C; centre; yellow nodes).

### Subjects with similar resting and task-general architectures demonstrate superior performance

We next examined whether cognitive performance and the efficiency of the transformation between rest and task-based neural architectures was related (Schultz and Cole, 2016). To this end, behavioural accuracies and update efficiencies were entered into linear regression analysis. Statistically significant (p < 0.05; FDR corrected for multiple comparisons) relationships between performance and update efficiency were identified in the three in-scanner tasks. These included relational reasoning (F(1,564) = 5.0, p = 0.026, r = 0.21), language processing (F(1,564) = 5.3, p = 0.021, r = 0.24), and working memory (F(1,564) = 4.9, p = 0.027, r = 0.20). Since metastability and update efficiency are correlated (in some connections) and since update efficiency predicts performance, we also confirmed that the efficiency-performance relations survived in the presence of controlling for global metastability. Overall, these findings suggests that cognitive ability is contingent upon resting state architecture being similar or ‘pre-configured’ to a task-general arrangement.

## Discussion

The present paper set out to answer a relatively simple question: is metastable neural dynamics higher at rest or during the performance of an explicit task? We sought to answer this question by comparing the metastability of the brain’s large-scale networks at rest and during the execution of several cognitively demanding tasks. Current theory suggests that spontaneous neural dynamics represent a repository of functional states from which more stable global brain states are constructed during task. Metastability between networks should therefore be maximal when subjects are at ‘cognitive rest’ and diminished during times of heightened cognitive demand. Surprisingly, our findings support an alternative possibility: metastability (or dynamic flexibility) between neural networks is actively enhanced by task engagement (Fig. 2). Explicit cognition was characterised by two types of network architecture: (1) a task-general network structure based on widespread changes in metastability; and (2) a task-specific network structure based on a small number of task-evoked changes in synchrony. Task-general architecture was principally characterised by high metastability in cognitive control networks and low metastability in sensory regions. Crucially, subjects with resting state architectures similar or ‘pre-configured’ to a task-general configuration displayed superior cognitive performance. Furthermore, although resting state networks were dynamically linked into context-dependent neurocognitive structures by task engagement, cognitive performance was more closely linked to the intrinsic activity of large-scale networks. High metastability in the intrinsic connectivity of cognitive control networks was associated with novel problem solving (or fluid intelligence) but was less important in tasks relying on previous experience (or crystallised intelligence). Overall, our findings suggest a key linkage between the intrinsic metastability of the brain’s large-scale network connectivity and cognition.

The present study used one measure of connectivity–metastability–to assess the dynamic stability within and between large-scale networks of the brain. Increased stability of dynamic functional connectivity appears to be a general property of cognitive engagement across multiple behavioural paradigms irrespective of the type of dynamic functional connectivity method employed (Cohen, 2018). Consistent with the properties of a critical system, dynamic functional connectivity is especially stable during tasks requiring sustained attention (Haimovici et al., 2013; Meisel et al., 2013). Focused cognition appears to induce the sub-critical dynamics necessary for reducing interference and optimising task performance (Fagerholm et al., 2015). Such dynamic stability is associated with increased integration across cognitive control networks and between cognitive control and other task-relevant networks (Elton and Gao, 2015; Hutchison and Morton, 2015; J. E. Chen et al., 2015). On the surface, increased stability between networks during task engagement appears at odds with the present finding of increased metastability. How can something be both flexible and stable at the same time? Fortunately, the paradox is resolved when we recognise that metastability and synchrony are correlated attributes of brain function (see Fig. S1 in the supplementary information). Thus, even though metastability (or the variation in synchrony) is increasing, so too is the average synchrony (which can be viewed as a measure of stability in this context). It should be emphasised that purely synchronous episodes of spatio-temporal coordination between networks is not reflective of typical neurocognitive processing. Our measure of synchrony reflects this reality by reporting the average synchrony (or mean phase coherence) over a period of time corresponding to a series of concatenated task blocks. Using this approach, we show that task blocks are characterised by on average higher levels of synchrony and higher levels of metastability at the same time. That is, synchrony (the mean phase coherence over time) and metastability (the variation in the mean phase coherence over time) increase together–a behaviour observed in cortical slice cultures excited pharmacologically (Yang et al., 2012). In light of this, synchrony and metastability should be considered as related rather than contrary methods of viewing brain activity. Two possibilities exist: (1) that metastability is the price paid for higher on average synchrony i.e. it is a form of noise; or (2) metastability is an adaptive process reflecting the ongoing engagement and disengagement of neural systems relevant to cognitive and affective states. Our finding, that cognitive performance is linked to the intrinsic metastability of large-scale networks connectivity, strongly supports a functional view of metastability. Indeed, theoretical and empirical findings suggest that intrinsic neural architecture is organised to support a range of functional states which can be a posteriori selected via exogenous input (Cabral et al., 2014; Hansen et al., 2014; Kringelbach et al., 2015; Ponce-Alvarez et al., 2015; Deco and Kringelbach, 2016; Deco et al., 2017). One may argue that metastability is being driven by the variation in synchrony obtained by artificially concatenating a series of disjoint task blocks. However, in principle, metastability within a single task block should also be enhanced on the provision that synchrony also increases. Presently, the temporal limits imposed by fMRI and the relatively short task runs preclude a direct test of this hypothesis.

Task engagement was principally characterised by enhanced metastability in the connectivity of cognitive control networks (Fig. 3). These included dorsal attention, fronto-parietal, cingulo-opercular, default mode, and ventral attention networks. The present finding is consistent with the notion that task-positive networks are areas of the brain that respond with activation increases to attention-demanding environments. The organisational state of the system may facilitate or inhibit the processing of incoming external stimuli by enacting a series of task-relevant synergies over the duration of the task (Kelso, 2009). Each of the networks presented similar but not identical patterns of engagement across the seven tasks. Two of the four, the dorsal attention and fronto-parietal networks, demonstrated consistent activity across all seven of the in-scanner tasks and were associated with the greatest increases in metastability. This raises the question of why dorsal fronto-parietal regions are involved in such a puzzling variety of tasks? One proposal is that cognitive computations relying on dorsal fronto-parietal areas are concerned with a single core function, namely ‘offline motor planning’ or ‘action emulation’ (Ptak et al., 2017). Such findings are consistent with the observation that hubs of the fronto-parietal network alter their pattern of functional connectivity with nodes of other networks based on goal-directed cognition in a highly adaptive domain-general manner (Cole et al., 2013, 2014b). Rhythmic attentional sampling linked to theta-band activity in the large-scale dorsal fronto-parietal regions (Fiebelkorn et al., 2018; Helfrich et al., 2018) may also account for increased metastability. Changes in metastability were not limited to interactions between cortical networks; they also involved subcortical, specifically, thalamo-cortical components. The finding of increased metastability between thalamus and dorsal fronto-parietal during task is consistent with empirical evidence showing that thalamic input wires the contextual associations upon which complex decisions depend into weakly connected cortical circuits (Halassa and Kastner, 2017; Schmitt et al., 2017).

The seven in-scanner tasks showed remarkably similar patterns of increased metastability between large-scale networks (Fig. 3). Such findings resonate with prior studies showing similar patterns of static functional connectivity between different tasks (Cole et al., 2014a; Schultz and Cole, 2016). Similarities between functional network configurations evoked under different behavioural paradigms constitute what has been referred to as a ‘task-general architecture’ (Cole et al., 2014a; Schultz and Cole, 2016). Our findings suggest that different behaviours recruit a similar set of network connections through metastable neural dynamics. We base this claim on three observations (1) increases in metastability were more widespread than increases in synchrony (Fig. 4); (2) increases in synchrony were highly specific to each task, whereas increases in metastability were more task-general (Fig. 5; Fig. 6); and (3) most of the variance (78%) between tasks was accounted for by a single principal component (Fig. 7). Our task-general configuration was organised into distinct regions of both high and low metastability (Fig. 7). Networks related to cognitive control exhibited dynamic flexibility whilst primary sensory networks favoured dynamic stability. These findings map onto our present understanding of cortical organisation and function. Primary sensory areas are responsible for processing a single modality whereas higher order association areas must integrate information into more complex representations e.g. language, executive function, attention, and memory. Thus, each step up the hierarchy entails integrating information from a greater diversity of sources, and this, in turn, is reflected in a region’s dynamic flexibility. From another perspective, widespread increases in metastability are quite surprising. They appear to indicate that most of the brain is involved during task which appears to contradict findings from cognitive fMRI studies showing only limited activation. One possibility is that some of these changes in metastability are linked to suppression of a given network during task performance.

Even during periods of apparent ‘rest’ a subject’s network connectivity continued to display strong integrative and segregative tendencies linked to cognitive performance. Remarkably, a subject’s ability to answer questions correctly, both in and out of the scanner, related to their intrinsic neural dynamics (Fig. 8). Curiously, task-based changes in metastability were not related to cognitive performance, rather, subjects whose dynamics explored a greater range of network configurations at rest demonstrated the highest cognitive test scores and behavioural accuracy. Unlike crystallised intelligence, which was largely unrelated to metastability, the flexibility of network states afforded by metastable neural dynamics was strongly linked to fluid intelligence. Accumulating evidence suggests that human intelligence arises from the dynamic reorganisation of brain networks (Barbey, 2018). Flexible network transitions may support the ‘difficult-to-reach’ networks states associated with novel problem solving but may be less relevant for accessing the ‘easy-to-reach’ network configurations associated with local knowledge and experience (Power and Petersen, 2013; Gu et al., 2015). The link between metastable neural dynamics and fluid intelligence was especially pronounced in systems involved in cognitive control. Our results are consistent with prior observations linking cognitive control capacity and fluid intelligence (Conway et al., 2002; Cole and Schneider, 2007; Cole et al., 2012). Such dynamic flexibility may be a fixed property of a subject’s neural architecture. We found that metastability at rest was commensurate with metastability during the execution of a task (see Fig. S3 in the supplementary information). Individual differences in metastable neural dynamics likely stem from a combination of innate and experiential influences on the anatomical connectivity between regions; a view consistent with prior observations linking reduced metastability to altered network topology (Hellyer et al., 2015; Váša et al., 2015; Córdova-Palomera et al., 2017; Alderson et al., 2018).

So why is metastability at rest predictive of behavioural and cognitive performance as opposed to metastability during the task itself? The idea that functional couplings between regions at rest contain information relevant to cognition, perception, and behaviour is supported by substantial empirical evidence (Sadaghiani, 2010; Sadaghiani and Kleinschmidt, 2013). Static resting state functional connectivity has been linked to a number of general cognitive abilities that include, amongst others, IQ, executive function, episodic memory and reading comprehension (for review, see Vaidya and Gordon, 2013). Thus, rather than simply reflecting invariant structural anatomy, historical co-activation patterns, or internal dynamics of local areas, intrinsic activity predicts subsequent perceptual processing (Hesselmann et al., 2008; van Dijk et al., 2008; Busch et al., 2009; Mathewson et al., 2009; Sadaghiani et al., 2009; Lou et al., 2014; Van Den Berg et al., 2016).

Cognitive performance was also linked to the efficiency of the transformation between rest and task-based network architectures. Specifically, subjects whose resting state was similar to the task-general architecture garnered the highest cognitive scores. Successful cognition is likely predicated on an adequate a priori dynamic configuration before the onset of task-relevant stimuli, as opposed to simple ad hoc adjustments after the fact (Bolt et al., 2018). Thus, resting state activity may reflect the brain’s predictive engagement with the environment (Sadaghiani, 2010; Sadaghiani and Kleinschmidt, 2013; Clark, 2016). Given that the resting state reflects previous experience and the anticipation of likely future events, a resting state network architecture ‘pre-configured’ to task is more in line with future cognitive requirements (Bar, 2011). Update efficiency, or the ability to switch from a rest- to a task-based configuration mapped onto our task-general architecture (Fig. 9). High update efficiency was associated with dynamic flexibility in dorsal attention and fronto-parietal control networks and dynamic stability in primary sensory networks. Such findings conform to our intuitive expectation that cortical networks require varying amounts of dynamic flexibility to fulfil their function. Presumably, sensory networks function even when subjects are cognitively at rest, hence, metastability increases less in these regions during times of heightened cognitive demand. Interestingly, several regions demonstrated high and low metastability components. These included the salience network of which the insula–a known site of multi-modal integration of sensory, motor, emotional, and cognitive information–is a part (Gogolla, 2017). Update efficiency based on a static measure of functional connectivity has been considered in a previous study (Schultz and Cole, 2016). Our work differs from this approach in that we consider the update efficiency within the context of a dynamic measure of network connectivity in which both integrative and segregative tendencies coexist (Kelso, 1995, 2012; Tognoli and Kelso, 2014b).

Potential limitations of the findings presented here should be noted. Firstly, it is worth emphasising that the division of neural activity into intrinsic and task-evoked activity may be an artificial distinction and not an actual division respected by neural properties (Bolt et al., 2018). Moreover, the finding of increased metastability during task relative to rest may, to some extent, be dependent on factors related to experimental design including: (1) the time frame considered; and (2) the temporal resolution of the imaging modality used. Increased metastability was identified in a series of concatenated task blocks. A more desirable experimental setup would compensate for fMRI’s lack of temporal precision by measuring metastability over an extended period within a single task block. Doing so would ameliorate the potential confound of introducing variations in synchrony– our definition of metastability–by including rest, cue, and fixation blocks. In the present paper, we mitigated this risk by estimating metastability with non-task block data removed. Moreover, due to restrictions on the length of the runs, metastability was estimated using the active and control components of the task, hence, participants were not exclusively performing the ‘cognitive task’ but the ‘control task’ designed to compensate for non-specific effects (e.g. the zero back condition in the working memory task). It is worth noting however, that in most cases, there was either no control condition (e.g. the motor task) or subjects still performed a meaningful exercise in the control task condition (e.g. the language task which comprised a story and math component). The results were robust in that metastability was not driven by bias associated with using time series of different lengths (we used the same number of data points for rest and task), nor by alternating periods of rest and task within a single run (we removed cue and fixation blocks). Despite attention to such factors, it remains of critical importance to establish the validity of observed synchronisation dynamics in light of recent findings questioning the link between time varying functional connectivity and task-relevant neural information (Hindriks et al., 2016; Laumann et al., 2017; Liégeois et al., 2017). Rapid changes in synchronisation may not be directly tied to external task demands, but rather to internally driven factors that include attention, motivation, arousal, fatigue, goals, or levels of consciousness (Kucyi, 2017; Kucyi et al., 2017). For instance, network integration during rest is linked with greater pupil diameter; a proxy for arousal, as well as better task performance (Shine et al., 2016a). Similarly, across repeated resting state scans of the same individual, higher levels of fatigue are related to more stable estimates of dynamic functional connectivity whereas higher levels of attention are related to more variable measures of dynamic functional connectivity (Shine et al., 2016b). Rapid dynamics likely include both meaningful neural information and physiological signals that relate to differences in rate or volume of blood flow, respiration, and heart rate (Cohen, 2018). Untangling the relationship between rapid dynamics, internal mentation, cognitive performance, affective states, physiological noise, arousal, fatigue, and general cognition will become increasingly important as we seek to understand the neural basis of human behaviour.

The present study used the theoretical framework of metastable coordination dynamics to examine how self-organising network interactions give rise to cognition (Kelso, 1995; Bressler and Kelso, 2001, 2016). The dual nature of this coordination–a simultaneous integration and segregation over space and time–occurred in the absence of input, suggesting that network interactions are not primarily reflexive or stimulus driven but spontaneously metastable (Tognoli and Kelso, 2014b). Crucially, intrinsic brain dynamics were linked to cognitive performance in a variety of behavioural domains, principally, in region associated with cognitive control. Moreover, subjects with resting state architectures similar or ‘pre-configured’ to their task-based configuration demonstrated the most accurate behaviour. Switching between rest- and task-based network architecture was contingent on high metastability in the connectivity of cognitive control networks and low metastability in the connectivity of primary sensory networks. Our analysis also revealed a critical linkage between key factors of general intelligence and endogenous brain activity, namely, high metastability in regions devoted to cognitive control during tests of fluid intelligence–a metric associated with novel problem solving, and low metastability in the same regions during tests of crystallised intelligence–a metric dependent on previous knowledge and experience. Overall, our findings suggest that cognitive function is attendant upon the brain’s intrinsic patterns of spontaneous neural activity.

In addition to spontaneous dynamics observed in the ‘resting’ brain, a second type of order was imposed externally in response to environmental and stimulus contingencies. Increased metastability during task is consistent with the dynamic reorganisation of resting state networks into large-scale neurocognitive entities (Bressler and Kelso, 2001, 2016). These variability reducing synergies likely constrain and exploit the enormous degrees of freedom afforded by the nervous system (Kelso, 2009). However, unlike spontaneous neural dynamics, the phenomenon of task-evoked metastability was unrelated to task performance.

Taken together, our findings suggest that the metastable regime of coordination dynamics offers considerable potential as a theoretical and conceptual framework for linking resting state network activity to cognition and behaviour. Although resting state networks are dynamically reconfigured during cognitive engagement, task onset impacts the spatiotemporal properties of a pre-existing functional architecture. Such coordinative behaviour, organised through spontaneous metastable dynamics, appears to contribute to cognition by anticipating incoming stimuli. Cognitive function is therefore not primarily stimulus-driven or reflexive but arises from the brain’s self-organising character.

## Acknowledgements

This work was supported by a Department for Employment and Learning Northern Ireland PhD studentship. This work was also supported, in part, by a grant given to ALWB from Science Foundation Ireland (grant number 11/RFP.1/NES/3194). JASK is supported by a grant from the National Institute of Mental Health (MH080838), the Chaire d’Excellence Pierre de Fermat, and the Davimos Family Endowment for Excellence in Science.

Data were provided [in part] by the Human Connectome Project, WU-Minn Consortium (Principal Investigators: David Van Essen and Kamil Ugurbil; 1U54MH091657) funded by the 16 NIH Institutes and Centers that support the NIH Blueprint for Neuroscience Research; and by the McDonnell Center for Systems Neuroscience at Washington University.

