## Supplementary Information for "Metastable neural dynamics underlies cognitive performance across multiple behavioural paradigms"

**Timing and trial structure**

Timing and trial structure are taken from:

<http://protocols.humanconnectome.org/HCP/3T/task-fMRI-protocol-details.html>

**Working memory**

The category specific representation task and the working memory task are combined into a single task paradigm. Participants were presented with blocks of trials that consisted of pictures of places, tools, faces and body parts (non-mutilated parts of bodies with no “nudity”). Within each run, the 4 different stimulus types were presented in separate blocks. Also, within each run, ½ of the blocks use a 2-back working memory task and ½ use a 0-back working memory task (as a working memory comparison). A 2.5 second cue indicates the task type (and target for 0-back) at the start of the block. Each of the two runs contains 8 task blocks (10 trials of 2.5 seconds each, for 25 seconds) and 4 fixation blocks (15 seconds). On each trial, the stimulus is presented for 2 seconds, followed by a 500 ms inter-task interval (ITI).

**Gambling**

This task was adapted from the one developed by Delgado and Fiez (Delgado et al., 2000). Participants play a card guessing game where they are asked to guess the number on a mystery card (represented by a “?”) in order to win or lose money. Participants are told that potential card numbers range from 1-9 and to indicate if they think the mystery card number is more or less than 5 by pressing one of two buttons on the response box. Feedback is the number on the card (generated by the program as a function of whether the trial was a reward, loss or neutral trial) and either: 1) a green up arrow with “$1” for reward trials, 2) a red down arrow next to -$0.50 for loss trials; or 3) the number 5 and a gray double headed arrow for neutral trials. The “?” is presented for up to 1500 ms (if the participant responds before 1500 ms, a fixation cross is displayed for the remaining time), following by feedback for 1000 ms.  There is a 1000 ms ITI with a “+” presented on the screen. The task is presented in blocks of 8 trials that are either mostly reward (6 reward trials pseudo randomly interleaved with either 1 neutral and 1 loss trial, 2 neutral trials, or 2 loss trials) or mostly loss  (6 loss trials pseudo-randomly interleaved with either 1 neutral and 1 reward trial, 2 neutral trials, or 2 reward trials).  In each of the two runs, there are 2 mostly reward and 2 mostly loss blocks, interleaved with 4 fixation blocks (15 seconds each).

**Motor**

This task was adapted from the one developed by Buckner and colleagues (Buckner et al., 2011; Yeo et al., 2011). Participants are presented with visual cues that ask them to either tap their left or right fingers, or squeeze their left or right toes, or move their tongue to map motor areas. Each block of a movement type lasted 12 seconds (10 movements), and is preceded by a 3 second cue. In each of the two runs, there are 13 blocks, with 2 of tongue movements, 4 of hand movements (2 right and 2 left), and 4 of foot movements (2 right and 2 left). In addition, there are 3 15-second fixation blocks per run. This task contains the following events, each of which is computed against the fixation baseline.

**Language processing**

This task was developed by Binder and colleagues (Binder et al., 2011) and uses the E-prime scripts provided by these investigators.  The task consists of two runs that each interleave 4 blocks of a story task and 4 blocks of a math task.  The lengths of the blocks vary (average of approximately 30 seconds), but the task was designed so that the math task blocks match the length of the story task blocks, with some additional math trials at the end of the task to complete the 3.8 minute run as needed.  The story blocks present participants with brief auditory stories (5-9 sentences) adapted from Aesop’s fables, followed by a 2-alternative forced-choice question that asks participants about the topic of the story.  The example provided in the original Binder paper (p. 1466) is “For example, after a story about an eagle that saves a man who had done him a favor, participants were asked, “Was that about revenge or reciprocity?” The math task also presents trials auditorially and requires subjects to complete addition and subtraction problems.  The trials present subjects with a series of arithmetic operations (e.g., “fourteen plus twelve”), followed by “equals” and then two choices (e.g., “twenty-nine or twenty-six”).  Participants push a button to select either the first or the second answer. The math task is adaptive to try to maintain a similar level of difficulty across participants.  For more details on the task, please see (Binder et al., 2011).

**Social cognition (theory of mind)**

Participants were presented with short video clips (20 seconds) of objects (squares, circles, triangles) that either interacted in some way, or moved randomly on the screen. These videos were developed by either Castelli and colleagues (Castelli et al., 2013) or Martin and colleagues (Wheatley et al., 2007). After each video clip, participants judge whether the objects had a mental interaction (an interaction that appears as if the shapes are taking into account each other’s feelings and thoughts), Not Sure, or No interaction (i.e., there is no obvious interaction between the shapes and the movement appears random). Each of the two task runs has 5 video blocks (2 Mental and 3 Random in one run, 3 Mental and 2 Random in the other run) and 5 fixation blocks (15 seconds each).

**Relational Processing**

This task was adapted from the one developed by Christoff and colleagues (Smith et al., 2007). The stimuli are 6 different shapes filled with 1 of 6 different textures.  In the relational processing condition, participants are presented with 2 pairs of objects, with one pair at the top of the screen and the other pair at the bottom of the screen.  They are told that they should first decide what dimension differs across the top pair of objects (differed in shape or differed in texture) and then they should decide whether the bottom pair of objects also differ along that same dimension (e.g., if the top pair differs in shape, does the bottom pair also differ in shape).  In the control matching condition, participants are shown two objects at the top of the screen and one object at the bottom of the screen, and a word in the middle of the screen (either “shape” or “texture”).  They are told to decide whether the bottom object matches either of the top two objects on that dimension (e.g., if the word is “shape”, is the bottom object the same shape as either of the top two objects.  For both conditions, the subject responds yes or no using one button or another.  For the relational condition, the stimuli are presented for 3500 ms, with a 500 ms ITI, and there are four trials per block.  In the matching condition, stimuli are presented for 2800 ms, with a 400 ms ITI, and there are 5 trials per block.  Each type of block (relational or matching) lasts a total of 18 seconds.  In each of the two runs of this task, there are 3 relational blocks, 3 matching blocks and 3 16-second fixation blocks.

**Emotion processing**

This task was adapted from the one developed by Hariri and colleagues (Hariri et al., 2002).  Participants are presented with blocks of trials that either ask them to decide which of two faces presented on the bottom of the screen match the face at the top of the screen, or which of two shapes presented at the bottom of the screen match the shape at the top of the screen.  The faces have either an angry or fearful expression.  Trials are presented in blocks of 6 trials of the same task (face or shape), with the stimulus presented for 2000 ms and a 1000 ms ITI.  Each block is preceded by a 3000 ms task cue (“shape” or “face”), so that each block is 21 seconds including the cue. Each of the two runs includes 3 face blocks and 3 shape blocks, with 8 seconds of fixation at the end of each run.

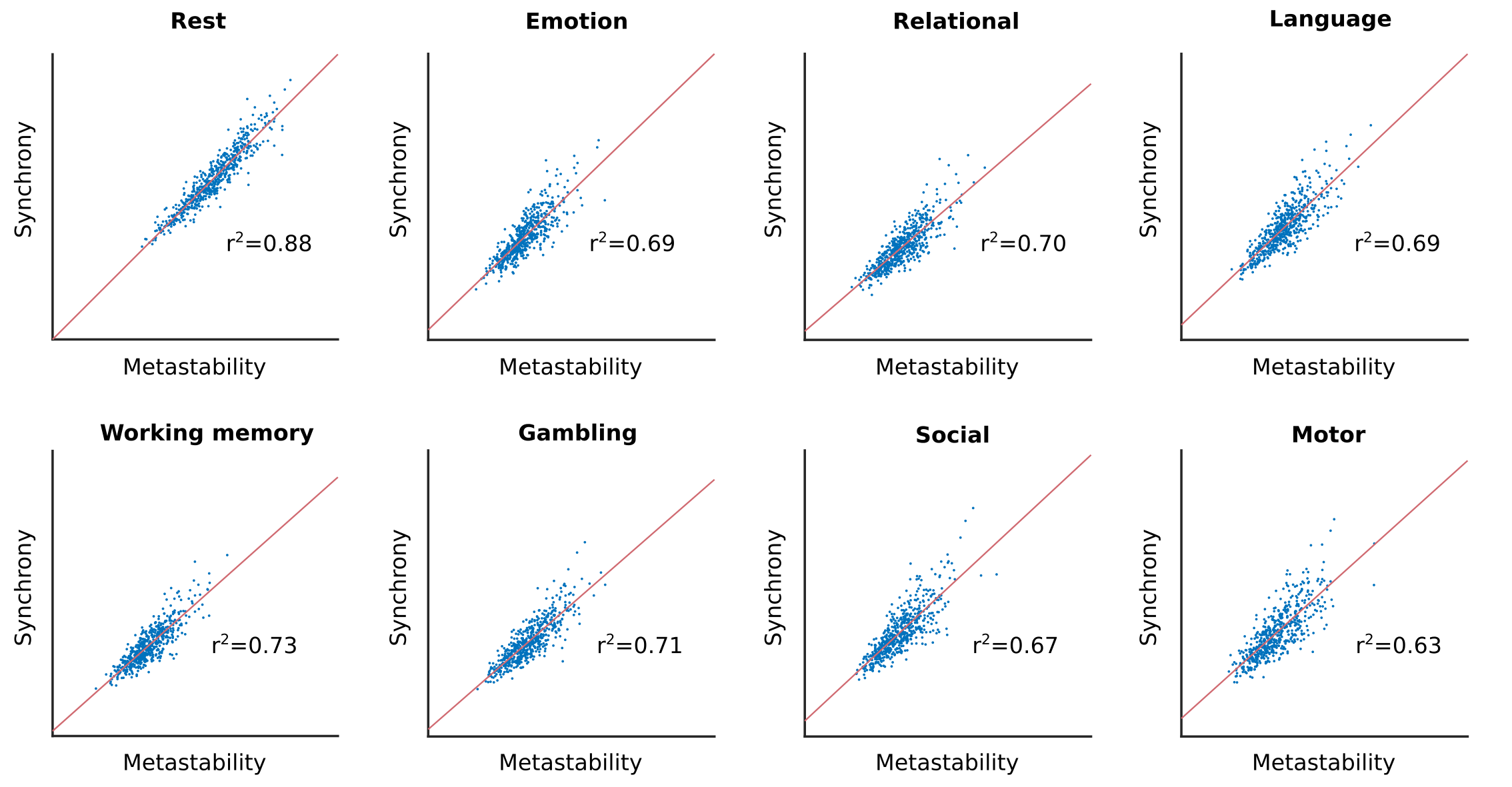

Figure 1: Synchrony and metastability are correlated. Global metastability versus global synchrony across multiple behavioural domains.

**Synchrony (mean phase coherence) and metastability (variation in mean phase coherence) are correlated quantities**

Synchrony plotted against metastability with line of best fit in red (Fig. 1). Across all seven tasks and rest, synchrony and metastability were strongly correlated quantities. Interestingly, synchrony and metastability were most strongly associated in the resting state. As the number of transitions between metastable brain states increases, so does the stability (or mean synchrony between brain regions).

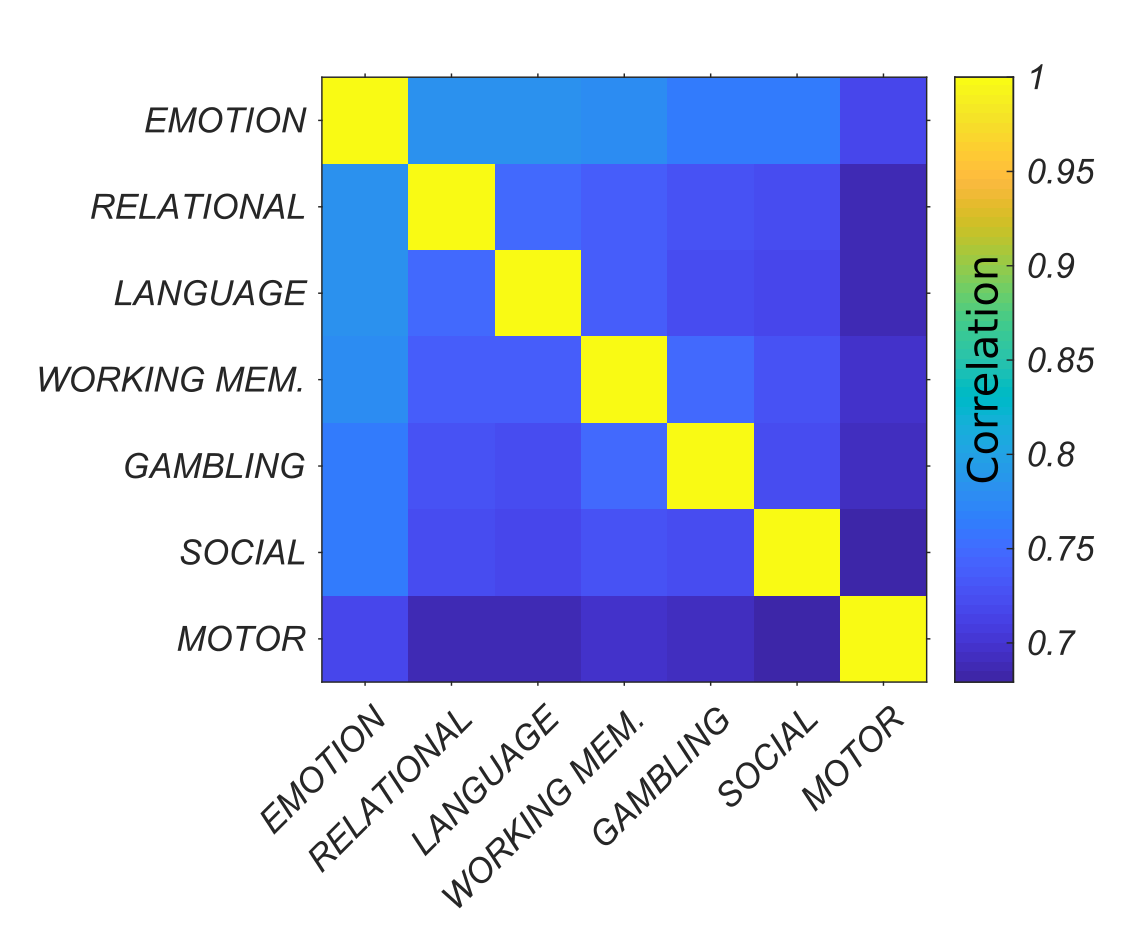

Figure 2: Different behaviours recruit a similar set of metastable interactions between large-scale networks. Interactions matrices (based on metastability) were correlated for each subject. The average for all subjects is reported.

**Highly similar task-based configurations are recruited by different behaviours**

Correlating each subject’s seven interaction matrices (based on metastable couplings between large-scale networks) and then taking the average for all subjects revealed that all seven tasks were attendant upon a similar network architecture.

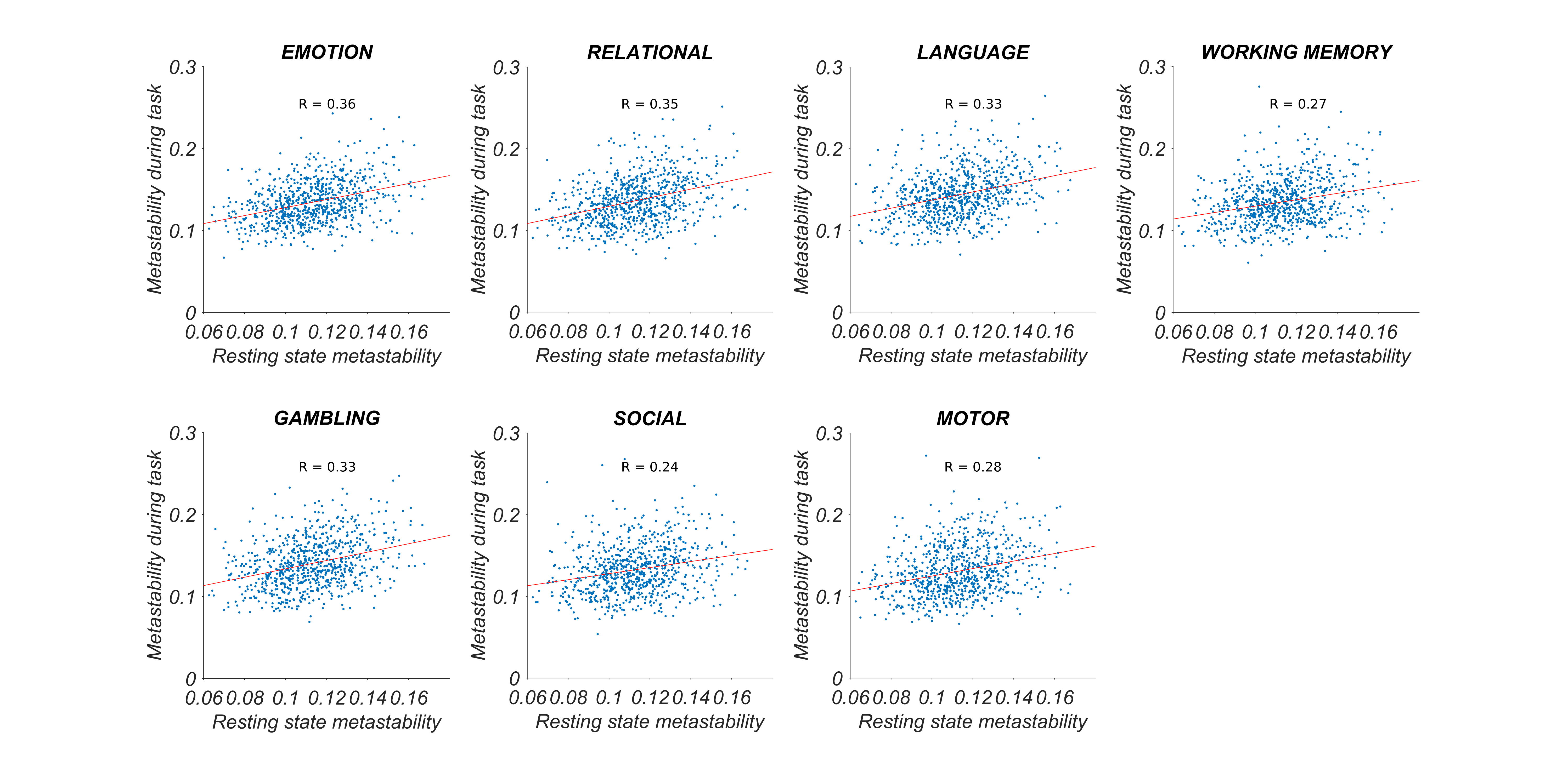

Figure 3: High metastability at rest is associated with high metastability during task performance. Statistically significant correlation between global metastability of fMRI BOLD signal at rest and global metastability during task performance (p < 0.001; corrected). Standardised effects sizes are indicated (Pearson’s r or correlation coefficient).

**High metastability at rest is commensurate with high metastability during task**

The fact that the metastability of individual network connections at rest is predictive of task performance raises an interesting question. Why is metastability at rest but not metastability during the execution of the task itself predictive of task performance? One hypothesis is that resting state neural dynamics captures the repertoire of functional couplings required to perform well under task-based conditions. Hence, a subject’s resting state metastability is an indication of their overall capacity for flexible network dynamics.

To test this hypothesis, we entered global task-based metastability (estimated from one of the seven in-scanner tasks) as dependent variable and global resting state metastability as predictor into a linear regression analysis. We found a significant positive correlation between global metastability at rest and global metastability during the execution of the seven in-scanner tasks (corrected for multiple comparisons at a p-level of 0.001). These included emotion perception (F(1,888) = 118, p < 0.001), relational reasoning (F(1,564) = 114, p < 0.001), language processing (F(1,564) = 97.2, p < 0.001), working memory (F(1,564) = 62.5, p < 0.001), gambling/reward learning (F(1,564) = 97.3, p < 0.001), social cognition/theory of mind (F(1,564) = 48.3, p < 0.001), and motor responses (F(1,564) = 69.3, p < 0.001). Taken together, these results suggest that task-based metastable neural dynamics are a stable property of a subject’s intrinsic neural architecture when at rest.
